# METTL7A and METTL7B are responsible for *S*-thiol methyl transferase activity in liver

**DOI:** 10.1101/2022.10.12.511968

**Authors:** Drake A. Russell, Marvin K. Chau, Yuanyuan Shi, Benjamin J. Maldonato, Rheem A. Totah

## Abstract

*S*-Methylation of drugs containing thiol-moieties often alters their activity and results in detoxification. Historically, scientists attributed methylation of exogenous aliphatic and phenolic thiols to a putative *S-*adenosyl-L-methionine dependent membrane-associated phase II enzyme known as thiol methyltransferase (TMT). TMT has a broad substrate specificity and methylates the thiol metabolite of spironolactone, mertansine, ziprasidone, captopril, and the active metabolites of the thienopyridine pro-drugs, clopidogrel, and prasugrel. Despite TMT’s role in the *S-*methylation of clinically relevant drugs, the enzyme(s) responsible for this activity remained unknown. We recently identified methyltransferase-like protein 7B (METTL7B) as an alkyl thiol-methyltransferase. METTL7B is an endoplasmic-reticulum-associated protein with similar biochemical properties and substrate specificity to TMT. Yet, the historic TMT inhibitor, 2,3-dichloro-α-methylbenzylamine (DCMB), has no effect on the activity of METTL7B, indicating that multiple enzymes contribute to TMT activity. Here we report that methyltransferase-like protein 7A (METTL7A), an uncharacterized member of the METTL7 family, also acts as a thiol-methyltransferase. METTL7A exhibits similar biochemical properties to TMT, including inhibition by DCMB (IC50 1.2 µM). Applying quantitative proteomics to human liver microsomes and gene modulation experiments in HepG2 and HeLa cells, we determined that TMT activity correlates closely with METTL7A and METTL7B protein levels. Furthermore, purification of a novel His-GST-tagged recombinant protein and subsequent activity experiments prove that METTL7A can selectively methylate exogenous thiol-containing substrates, including 7α-thiospironolactone, dithiothreitol, 4-chlorothiophenol, and mertansine. We conclude that the METTL7 family encodes for two enzymes, METTL7A and METTL7B, which we have renamed TMT1A1 and TMT1B1, respectively, that are responsible for TMT activity in liver microsomes.

**Significance Statement:** We identified METTL7A (TMT1A1) and METTL7B (TMT1B1) as the enzymes responsible for the microsomal alkyl thiol methyltransferase activity. These are the first two enzymes directly associated with microsomal TMT activity. *S-*Methylation of many commonly prescribed thiol-containing drugs alters their pharmacological activity and/or toxicity, and identifying the enzymes responsible, will improve our understanding of the DMPK properties of alkyl- or phenolic-thiol-containing therapeutics.

**Visual Abstract:** 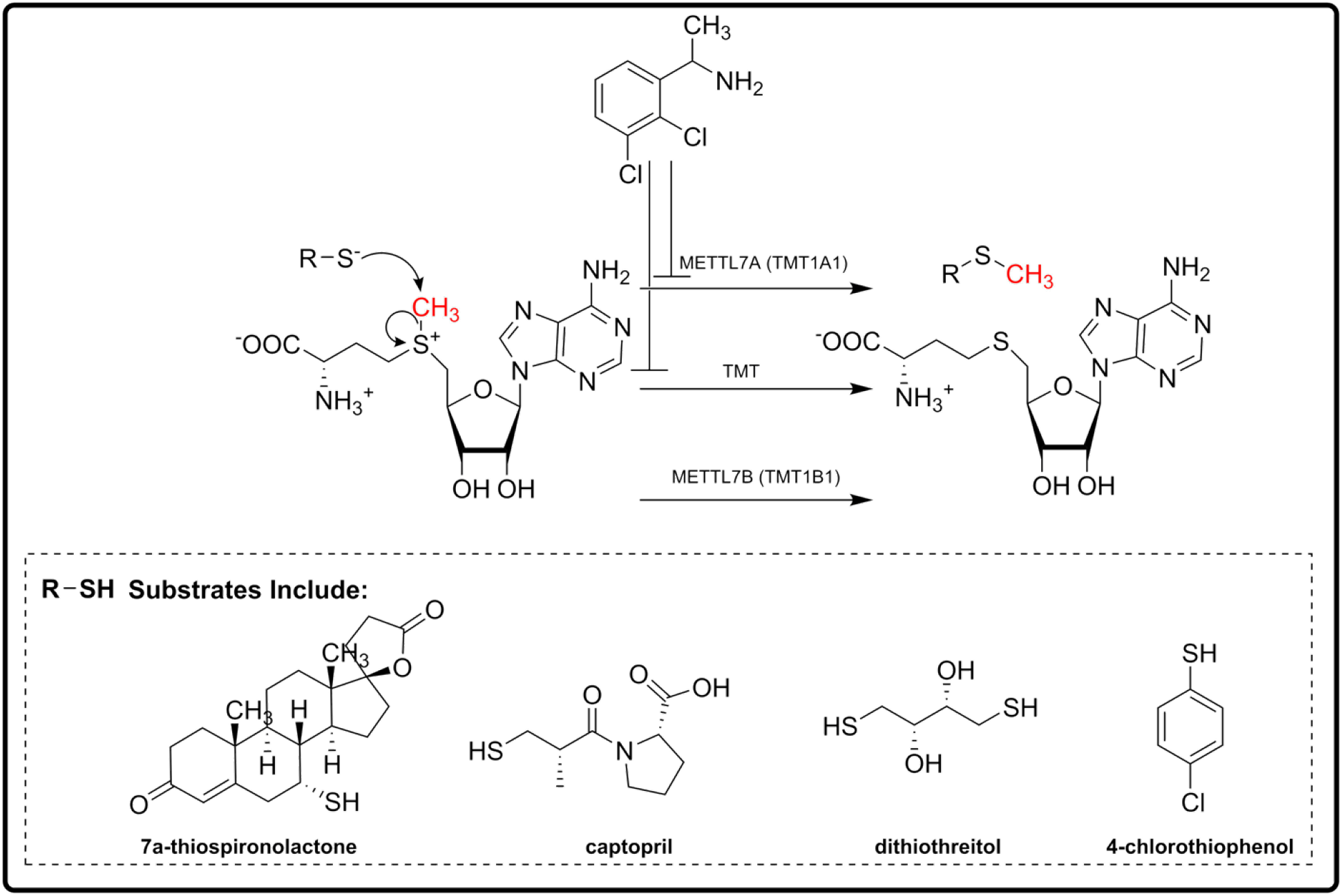

## Introduction

In humans, there are two distinct enzymes with *S-*methyltransferase activity, thiopurine methyltransferase (TPMT) and thiol methyltransferase (TMT) (Bremer and Greenberg, 1961; Yates et al., 1997). Both TPMT and TMT transfer a methyl group from the cofactor *S*-adenosyl-L-methionine (SAM) to a thiol moiety. Thiols are reactive nucleophiles capable of altering the oxidation potential of essential biomolecules by forming damaging covalent bonds. TPMT is a well-characterized methyltransferase that specifically methylates exogenous thiopurines (Krynetski and Evans, 2003). TPMT substrates such as 6-mercaptopurine and azathioprine are a class of chemotherapeutic agents and immunosuppressive drugs with a narrow therapeutic index, and *S-*methylation leads to their inactivation. Genetic polymorphisms in TPMT that result in decreased methylation activity can lead to toxicity from these drugs (Krynetski and Evans, 2003; Wusk et al., 2004). Weinshilboum and Sladek identified the enzyme responsible for TPMT activity in 1980 (Weinshilboum and Sladek, 1980), and many studies detailing the biochemical and structural characterization of TPMT have since followed. On the other hand, the enzymes involved in TMT activity were not known and as a result, historically, the microsomal fractions of various tissue homogenates were used to characterize TMT activity. Past research established that TMT does not share any substrates with TPMT but is responsible for the SAM-dependent *S*-methylation of exogenous alkyl and phenolic thiol-containing compounds. Endogenous thiols such as glutathione and cysteine are not substrates for TMT.

Many of the drugs that TMT methylates require their thiol functional group for activity, and methylation can dramatically alter efficacy. Potential TMT substrates include 7α-thiospironolactone (TSL), mertansine, the active metabolites of prasugrel and clopidogrel, ziprasidone, and captopril (Keith et al., 1984; Erickson et al., 2010; Obach et al., 2012; Kazui et al., 2014; Liu et al., 2015; Taplin et al., 2018). In most cases, it is difficult to determine *a priori* how methylation will alter the activity of a particular drug. For example, the *S*-methyl metabolite of spironolactone, 7α-thiomethylspironolactone (TMSL), is the major circulating metabolite and is responsible for most of spironolactone’s activity in vivo (Overdiek and Merkus, 1987). Mertansine, a potent microtubule inhibitor that is conjugated to antibodies by a disulfide bond, contains a free thiol upon reduction from the attached antibody and incorporation into a cell (Bargh et al., 2019). Interestingly, one report suggested that *S*-methylation of mertansine’s free thiol alters its interaction with microtubules by increasing mertansine’s effect on the dynamic instability of microtubules and promoting their aggregation; this is a widely different mechanism of action compared to the parent, free thiol, which inhibits microtubule polymerization (Lopus et al., 2010). Clopidogrel and prasugrel are thienopyridine pro-drugs that require activation via P450-mediated ring opening to expose a reactive thiol that directly adducts and irreversibly inhibits P2Y12, a transmembrane receptor that controls platelet aggregation (Savi et al., 2006). *S*-methylation of the reactive thiol results in the inactivation of these agents due to their inability to covalently link to their receptor (Savi et al., 2006). A better characterization of the enzymes responsible for TMT activity would provide clinicians and drug developers alike, much needed insight when faced with a compound that is metabolized by TMT.

Recently, we identified methyltransferase-like protein 7B (METTL7B) as an enzyme capable of catalyzing SAM-dependent methylation of alkyl thiols (Maldonato et al., 2021). METTL7B is a 28 kDa microsomal protein with a similar tissue distribution and substrate specificity to TMT (Zehmer et al., 2008). Although there is no specific inhibitor for TMT, 2,3-dichloro-α-methylbenzylamine (DCMB) potently inhibits TMT and has been used to confirm TMT activity in liver microsomes (Glauser et al., 1993; Maw et al., 2018). Interestingly, despite its similarities to TMT, DCMB does not inhibit METTL7B and thus we hypothesized that additional enzymes may account for microsomal TMT activity.

The gene “*putative methyltransferase-like protein 7A* (*METTL7A*)” codes for the protein METTL7A that belongs to the same family as METTL7B (Zehmer et al., 2008). METTL7A is proposed to be a membrane-associated methyltransferase and shares 75% sequence homology with METTL7B, but biochemical characterization that defines its endogenous function is scant. In this work, we report that METTL7A, like METTL7B, is an *S*-methyltransferase with many similarities to the historic TMT, including potent inhibition by DCMB. Using multiple approaches including quantitative proteomics, single donor human liver microsomes (HLM), and gene modulation experiments in HepG2 and HeLa cells, we determine that TMT activity is strongly associated with METTL7A protein levels. Furthermore, the purification of a novel His-GST-tagged recombinant protein and extensive biochemical characterization established that METTL7A methylates a wide variety of thiol-containing substrates, including TSL, dithiothreitol (DTT), 4-chlorothiophenol, and mertansine.

## Materials and Methods

### Chemicals, Materials, and Reagents

<u>Activity assay substrates, reagents, cofactor, inhibitor, and internal standards:</u> phenylethanolamine, dopamine, cysteine, 4-chlorothiophenol, and L-penicillamine from Sigma Aldrich (St. Louis, MO); mertansine and 1,2-Dimyristoyl-sn-glycero-3-PG (DMPG) were obtained from Cayman Chemicals (Ann Arbor, Mi); 7α-thiospironolactone was produced in-house from spironolactone using a previously published method (Agusti et al., 2013); spironolactone, reduced glutathione, captopril, and DL-1,4-dithiothreitol were obtained from Fisher Scientific (Hampton, NH); *S*-methyl mertansine, and 7α-thiomethylspironolactone-d7 was obtained from Toronto Research Chemicals (Toronto, ON, Canada); *S-*adenosyl-L-methionine was obtained from New England Biolabs (Ipswich, MA); and 2,3-dichloro-alpha-methylbenzylamine was sourced from Enamine (Monmouth Jct., NJ). <u>Reagents and materials for</u> *<u>E. coli</u>* <u>culture, protein purification, and activity assay experiments:</u> ampicillin sodium salt, isopropyl-β-D-thiogalactopyranoside, yeast extract, tryptone, NaCl, KH2PO4, K2HPO4, Thermo Scientific™ Halt™ Protease Inhibitor Cocktail, glycerol, CHAPS, white 96-well microplate, Nunc™ 96-Well Polypropylene Storage Microplates, Nunc™ 96-Well Slit-Cap Mats, and ultra-15 10 K molecular weight cut off centrifugal filter units were obtained from Fisher Scientific (Hampton, NH); His60 Ni Superflow Resin from Takara (Mountain View, CA); 5mL GSTrap Fast Flow column from Cytiva (Marlborough, MA); deoxyribonuclease I, lysozyme, and imidazole were obtained from Sigma Aldrich (St. Louis, MO); and MTase-Glo™ Methyltransferase Assay Kit was obtained from Promega (Madison, WI). All primers that are described for the sequencing or production of the N-GST METTL7A plasmid were obtained from Integrated DNA Technologies (San Diego, Ca). <u>Reagents and materials for activity assay cleanup and downstream LC/MS analysis:</u> acetonitrile (LC/MS grade), formic acid (LC/MS grade), acetic acid (LC/MS grade), Optima water, barium hydroxide, and zinc sulfate were obtained from Fisher Scientific (Hampton, NH); maleimide was obtained from Sigma Aldrich (St. Louis, MO). <u>Reagents and materials for SDS-page and western blot analysis:</u> c, NuPAGE™ MOPS SDS Running Buffer (20X), PageRuler Plus Prestained Protein Ladder (10 to 250 kDa), GelCode™ Blue Safe Protein Stain, iBlot™ Nitrocellulose Transfer Stacks, and METTL7B Rabbit anti-Human Polyclonal antibody were sourced from Fisher Scientific (Hampton, NH); Polyclonal Rabbit anti-Human METTL7A Antibody was obtained from LifeSpan BioSciences (Seattle, Wa); DYKDDDDK (Anti-FLAG) Tag Rabbit monoclonal antibody, β-actin rabbit monoclonal antibody, and GST-Tag rabbit monoclonal antibody were obtained from Cell Signaling (Danvers, MA); IRDye 680RD Goat anti-Rabbit secondary antibody was obtained from LiCor (Lincoln, NE); and Rabbit Polyclonal Anti-METTL7A Antibody was obtained from Origene (Rockville, MD). <u>Reagents and materials for cell culture and gene modulation:</u> Eagle’s Minimum Essential Medium, Gibco™ Opti-MEM™, Gibco™ Penicillin-Streptomycin, Corning™ Falcon™ Polystyrene 12-well and 6-well Microplates, Gibco™ Trypsin-EDTA (0.25%), Cytiva HyClone™ Fetal Bovine Serum, Invitrogen™ Lipofectamine™ 3000 & P3000 Transfection Reagents, and cell culture treated T75 plates were obtained from Fisher Scientific (Hampton, NH); Empty-, METTL7A-, and METTL7B-pCMV6 expression vectors were sourced from Origene (Rockville, MD); Dulbecco′s Phosphate Buffered Saline, METTL7A-specific siRNA, METTL7B-specific siRNA, and scramble non-specific negative control siRNA were obtained from Sigma Aldrich (St. Louis, MO). <u>Reagents for gene expression analysis:</u> Invitrogen™ MagMAX™ Total RNA Isolation Kit, Applied Biosystems™ High-Capacity RNA-to-cDNA™ Kit, Applied Biosystems™ TaqMan™ Fast Advanced Master Mix, and RT-PCR primers specific for METTL7A, METTL7B and GUSB were obtained from Fisher Scientific (Hampton, NH).

### METTL7A Expression and Purification

Recombinant wild-type METTL7A (N-GST-METTL7A) was cloned in *E. coli* using a unique expression plasmid that was created in-house. The expression plasmid backbone (pET21-10XHis-GST-HRV-dL5) was a gift from Marcel Bruchez (Addgene plasmid # 73214; http://n2t.net/addgene:73214; RRID: Addgene_73214). An open reading frame sequence that codes for METTL7A (Uniprot: Q9H8H3) was synthesized by BlueHeronBio (Bothell, WA) and inserted into the plasmid using BamHI and EcoRI restriction sites and general molecular biology techniques. The forward primer for insertion of the METTL7A sequence is as follows: 5′ CTAGCTAGGGATCCGCTCCGGCACCGGCTCCGGCACCGGCACCGATGGAGTTGACTATCTTCATCTTGCGCCTG 3′. The reverse primer for insertion of the METTL7A sequence was 5′ CTAGCTAGGAATTCTTATTAACGCGTTTTGACCGCGTA 3’. All plasmid inserts were validated by sequencing through Eurofins Genomics (Louisville, KY) and the sequencing histograms were analyzed using FinchTV software. The forward sequencing primer was 5′ GGGCTGGCAAGCCACGTTTGGTG 3′, and the reverse sequencing primer was 5′ CGTACCACTTCCACAAGTAAGTA 3′. The final expression plasmids positioned an N-terminal His-GST affinity tag onto the wild-type METTL7A protein sequence separated by a 10-residue poly-alanine-proline (AP)5 linker.

Expression plasmids were propagated using heat-shocked Stellar cells (Takara, Mountain View, CA, Ref: 636763, Lot: 2106487A). Competent LOBSTR-BL21(DE3) *E. coli* (Kerafast, Winston-Salem, NC, Ref: EC1002) transformed via heat shock with plasmids validated by sequencing. *E. coli* cells were cultured in an orbital shaker at 250 rpm, 37 °C, and in the presence of 100 µg/mL ampicillin.

To express recombinant protein in LOBSTR-BL21(DE3) cells, overnight cultures were added to ampicillin-containing TB expression media at a ratio of 1:100. Cells were grown for 4 h under normal growth conditions. METTL7A production was initiated via the addition of isopropyl β-D-1-thiogalactopyranoside (IPTG) to a final concentration of 1 mM. The temperature was reduced to 15 °C, and the cells were grown for an additional 24 h. Cells were harvested via centrifugation at 2500 RCF for 30 min, and the resulting pellets were either lysed immediately or stored at – 80 °C until future processing.

Frozen cell pellets were thawed on ice in a 4 °C cold cabinet overnight prior to resuspension in lysis buffer (50 mM KPi pH 7.0, 20% glycerol, 150 mM NaCl, 10 mM CHAPS, EDTA-free Halt Protease Inhibitor Cocktail) supplemented with 100 µg/mL lysozyme (Sigma Aldrich, St. Louis, MO, Ref: L-6876, Lot: 65H7025). The lysate was placed on a rocker at 4 °C until it became extremely viscous, then 100 µg/mL DNA Nuclease I (Sigma Aldrich, St. Louis, MO, Ref: DN25-100MG, Lot: SLCB6648) was added and the lysate was placed on a rocker at 4 °C until it was no longer viscous. The lysate was then centrifuged at 48,000 x g for 30 min at 4 °C, and the resulting supernatant was retained for subsequent purification steps.

The purification was performed using a vertical column hand-packed with 4 mL of His60 Ni Superflow Resin, followed by GST column purification using the ÄKTA start chromatography system (GE Healthcare, Chicago, IL). Cell lysate supernatant was added to the column containing nickel resin and it was recirculated across the column by a peristaltic pump overnight at 0.5 ml/min. The column was washed with 50 column volumes of His60 Ni purification buffer (50 mM KPi pH 7.0, 20% glycerol, 10 mM CHAPS, 300 mM NaCl) containing 50 mM imidazole. The protein was eluted from the column with 10 column volumes of His60 Ni purification buffer containing 300 mM imidazole.

The HisPur Ni–NTA column eluent was directly applied to a pre-conditioned GSTrapFF column at a flow rate of 1 mL/min and recirculated for 4 h. The column was then washed with GSTrapFF purification buffer (50 mM KPi pH 7.0, 20% glycerol, 10 mM CHAPS, 150 mM NaCl) until the absorbance at 280 nm decreased to baseline. The recombinant protein was eluted from the column using purification buffer, which contained 10 mM reduced glutathione and was adjusted to pH 8.0. The eluent was concentrated using MilliporeSigma™ Amicon™ Ultra-15 10 K molecular weight cut-off centrifugal filter units and the final protein concentration was determined by A280 measurement. Purified protein stocks were aliquoted and stored at – 80 °C for future use.

### Protein purity analysis

All SDS-PAGE Coomassie stained or western blot analysis were conducted with Invitrogen™ NuPAGE™ 4 to 12%, Bis-Tris, 1.0 mm, Mini Protein Gels in the XCell SureLock Mini-Cell Electrophoresis system using PageRuler Plus Prestained Protein Ladder as a molecular weight marker. Gels were run at room temperature and a constant 200 V. Total protein purity was determined with GelCode™ Blue Stain Reagent, a colloidal Coomassie dye. For western blot analysis, proteins were transferred to nitrocellulose membranes using the iBlot™ Gel Transfer Device. Primary antibody incubations were conducted overnight at 4 °C at a 1:500 dilution with rabbit anti-METTL7A (LifeSpan BioSciences, Seattle Wa, Ref: LS-C82875, Lot: 178089), anti-METTL7B (Invitrogen, Carlsbad, CA, Ref: PA5-58478, Lot: XC3518324A), or anti-GST (Cell Signaling, Danvers, MA, #2625S, Lot:8) primary antibodies. The secondary incubation was performed at a 1:10,000 dilution at room temperature for 1 h with IRDye 680RD goat anti-rabbit secondary antibody (LiCor, Lincoln, NE, Ref: 926-68071, Lot: D11102-15). Western blots and Coomassie-stained gels were visualized with an Odyssey gel scanner and blot images were analyzed using Image Studio Version 4.0 software.

### In vitro 7α-thiospironolactone and captopril methylation using recombinant METTL7A and METTL7B

For methyl transferase activity assays, aliquots of frozen purified recombinant METTL7A or METTL7B were diluted with reaction buffer (50 mM KPi pH 7.0, 20% glycerol, 150 mM NaCl, 10 mM CHAPS) to 0.2 mg/mL. The diluted protein was then combined with two parts reaction buffer containing 9 mg/mL DMPG. A fraction of diluted protein was boiled for 10 min while the remaining active protein was incubated on ice for 30 min to equilibrate with DMPG.

For the TSL methylation assay, TSL was dissolved in acetonitrile and then deposited in the appropriate wells of a 96-well reaction plate. The plate was dried under a vacuum for 10 min to remove the organic solvent. The diluted recombinant protein was then added to the plate and placed on a shaker at room temperature and 625 RPM for 5 min. DCMB, dissolved in water, was added to wells requiring DCMB and an equal volume of water was added to the remaining wells as a vehicle control. The reaction was initiated by adding SAM to a final concentration of 100 µM. The final TSL concentration was 50 µM, the final DCMB concentration was 100 µM, and the final protein concentration was 0.07 mg/mL. The plate was again placed on a plate shaker, sealed with a silicon plate mat, and then incubated at 37 °C for 30 min. For IC50 determination, METTL7A was incubated with 50 µM TSL, DCMB was added at 100 µM, 50 µM, 10 µM, 1 µM, 500 nM, 100 nM, 10 nM, and 1 nM final concentration.

After 30 min, the reaction was quenched with 5 × the reaction volume with ice-cold ethyl acetate containing 5 µM 7α-thiomethylspironolactone-d7 deuterated internal standard (TMSL-D7). During the quench step, each sample was mixed with ethyl acetate by gently pipetting up and down. The plate was then immediately frozen at −80 °C for 12 min. After 12 min, the aqueous layer was frozen, and the liquid organic top layer was transferred to a new plate. The transferred samples were dried with gentle heat and nitrogen gas flow (Biotage, Charlotte, NC) followed by reconstitution with 50:50 MeCN:H2O containing 0.2% acetic acid. Samples were analyzed with a Waters Xevo TQS mass spectrometer paired with a Waters Acquity LC using a 2.1 × 50 mm Waters 1.7 µm C18 column and 0.2% acetic acid in water and 0.2% acetic acid in MeCN as solvents A and B respectively. Consistent elution of TMSL and TMSL-D7 was achieved with an isocratic gradient held at a 50:50 ratio of solvent A to solvent B. Flow rate was constant at 0.4 mL/min. The transitions used to monitor for methyl metabolite 7α-thiomethylspironolactone (TMSL) and IS TMSL-D7 were 389.223>341.308 and 396>347.05, respectively, and TSL methylation was reported as the peak area ratio of TMSL / TMSL-D7.

For captopril methylation assay, captopril, dissolved in water, was added to 96-well plate wells containing diluted active or boiled protein, prepared as described above. A no enzyme control incubation was prepared by replacing the protein-containing portion of the components with reaction buffer. DCMB dissolved in water, or water as a vehicle control, was added to the appropriate wells. The final protein concentration was 0.07 mg/mL; for wells including DCMB, the final DCMB concentration was 100 µM. The plate was placed on a shaker for 2 min at 625 RPM at room temperature and the reaction was initiated by the addition of SAM to a final concentration of 100 µM. After initiating the reaction, the plate was mixed on the shaker, sealed with a silicon plate mat, and incubated at 37 °C for 25 min. The reaction was quenched by adding 15% zinc sulfate (w/v) at 1/8th the reaction volume. The quenched solution was incubated on ice for 10 minutes before a solution of saturated barium hydroxide, containing the d3-*S-* methyl captopril internal standard, was added at a 1:1 v/v with zinc sulfate. The plate was mixed at 625 RPM for 2 min and then incubated on ice for another 10 min. The 96-plate was centrifuged at 4,500 × g for 15 min to pellet all precipitated proteins and salts. The supernatant was transferred to a new 96-well plate and NaOH was added a final concentration of 125 mM. Finally, maleimide was added to the reaction to a final concentration of 500 mM. Maleimide reacts with unmethylated captopril to remove excess captopril from downstream analysis. The plate was sealed with a silicon plate mat, wrapped in foil, and incubated at room temperature for 1 hour. The plate was centrifuged again to precipitate any salts that formed during the alkylation reaction, the supernatant was transferred to a new 96-well plate, and was analyzed by LC-MS/MS. LC-MS/MS conditions for monitoring captopril methylation were similar to those described previously by Maldonato et al. (Maldonato et al., 2021). Captopril methylation was reported as the peak area ratio of Me-captopril / D3-Me-captopril subtracted from the average peak area ratio determined for the no-enzyme control group (the background).

### In vitro Michaelis-Menten kinetics using recombinant METTL7A and METTL7B

Purified N-GST-METTL7A, or N-GST-METTL7B, was diluted into a DMPG-containing buffer, and TSL was deposited into the wells of a 96-well plate as described above.

For single time point experiments with increasing concentrations of SAM, TSL was added to a final concentration of 250 µM, and SAM was added at 1 mM and 250, 100, 50, 25, 10, 5, and 1 µM to initiate the reaction.

For the SAH and SAM single time point activity assay matrix, TSL was added to a final concentration of 250 µM, SAH was added at 100, 50, 10, 5, and 0 µM, and SAM was added at 1 mM, 250, 100, 50, 25, 10, 5, and 1 µM to initiate the reaction.

For the TSL single time point activity assay with increasing concentrations of TSL, TSL was added to a final concentration of 1 mM, 250, 100, 50, 25, 10, 5, and 1 µM. SAM was added to a final concentration of 100 µM to initiate the reaction.

All reactions were incubated for 30 min at 37 °C before quenching with ethyl acetate containing TMSL-D7. The reactions were quenched, processed, and analyzed similar to the procedure described above.

### Substrate Screening

Potential substrates were screened using the Promega MTase-Glo™ Methyltransferase Assay Kit. Purified N-GST-METTL7A was diluted into a DMPG-containing buffer similar to the procedure described above. The substrate screening assay was performed in a 96-well plate at a final protein concentration of 0.07 mg/mL. Each probe substrate (1 mM final concentration) was added to the diluted enzyme. Samples were pre-equilibrated with shaking at 625 RPM for 5 min at room temp, and the reactions were initiated by adding SAM (100 µM). The plate was sealed with a silicon plate mat and incubated at 37 °C for 1 h. Enzymatic activity was quenched by adding DCMB to a final concentration of 100 µM. A volume from each well in the enzyme reaction plate was transferred to a new well in a white-walled 96-well flat bottom plate and then the plate was processed according to the manufacturer’s protocol. Luminescence was recorded from each well using a Synergy HTX Multi-Mode Reader (BioTek, Winooski, VT). All compounds were tested in triplicate with boiled and active N-GST-METTL7A, and relative turnover was determined by subtracting the response of boiled N-GST-METTL7A from active N-GST-METTL7A for each compound. The resulting difference for a no substrate control was then subtracted from all the active minus boiled differences that were determined for each compound using the following equation:

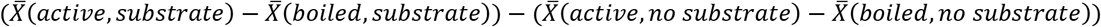

We noticed a significantly high background signal when using the Promega MTase-Glo™ Methyltransferase Assay Kit which is likely due to the presence of a thiol compound in the developing kit that is readily methylated by METTL7A or METTL7B. It is necessary to be cautious in designing controls, quenching the reaction or interpreting the data.

### Silencing METTL7A and METTL7B genes in HepG2 Cells

HepG2 cells (ATCC, Manassas Va, Ref: HB-8065) were cultured in Eagle’s minimum essential media (EMEM) with 10% fetal bovine serum, 0.1% v/v penicillin-streptomycin and seeded into 12-well tissue culture plates at 200K cells/well (n=3). Cells were reverse transfected with Lipofectamine 3000 and siRNA by following the manufacturer’s protocol for gene knockdown. The final concentration of small interfering RNA (siRNA) per treatment was 10 nM, and the final concentration of lipofectamine 3000 reagent was 2 µL/mL. Cells were reverse transfected with siRNA targeting *METTL7A, METTL7B, METTL7A* and *METTL7B* (combo), or scramble non-specific siRNA (Sigma Aldrich St. Louis, MO, Ref: METTL7A/EHU153411, METTL7/EHU008171, Scramble/SIC001). Cells were incubated with siRNA for 48 hours, washed with warm DPBS, and treated with serum-free EMEM containing TMT substrates (125 µM TSL or 500 µM captopril) in the presence or absence of 20 µM DCMB. Cells were incubated with TMT substrates for 24 hours after which the media was collected and analyzed for methylated metabolites by LC-MS/MS. The LC-MS/MS analysis of captopril and TSL were conducted as described above. The results were normalized to the media collected from negative control scramble siRNA-treated cells. The knockdown efficiency analysis were conducted as described by Maldonato et al., 2021 using proprietary TaqMan FAM reporter primers for *METTL7A, METTL7B*, and the housekeeping gene *GUSB*.

### Overexpression of METTL7A and METTL7B in HeLa Cells

HeLa cells (ATCC, Manassas Va, Ref: CCL-2™) were cultured in EMEM with 10% fetal bovine serum, 0.1% v/v penicillin-streptomycin and seeded into 6-well tissue culture plates at 500K cells/well (n=3). The cells were reverse transfected using Lipofectamine 3000 and P3000 reagent along with different pCMV expression vectors following the manufacturer’s protocol. The final concentration of pCMV plasmid expression vector per treatment was 1250 ng/mL, the final concentration of Lipofectamine 3000 reagent was 1.5 µL/mL, and the final concentration of P3000 reagent was 1 µL/mL. Cells were incubated with media containing Lipofectamine reagents and either a METTL7A (Origene, Rockville MD, Ref: RC202601), METTL7B (Origene, Rockville MD, Ref: RC203838), or empty control (Origene, Rockville MD, Ref: PS100001) expression vector for 36 h. After 36 h, the cell media was removed, the cells washed with warm DPBS, and the media replaced with serum-free EMEM containing 125 µM TSL +/-20 µM DCMB. Cells were incubated with TMT substrate for 24 h before the media was collected and analyzed for methylated metabolites by LC-MS/MS. The LC-MS/MS analysis of TSL was conducted as described above.

Protein overexpression was confirmed by western blot analysis. After incubation with the expression vector for 36 hours, cells were harvested by scraping each well with 1 mL of DPBS. Cells were then transferred to a 1.5 mL Eppendorf tube and pelleted by centrifuging for 5 min at 750 × g, 4 °C. The DPBS was removed and the cells from each treatment were lysed in 150 µL of RIPA buffer containing 1X halt protease inhibitor cocktail. The lysis was performed by resuspending cells in lysis buffer and pipetting up and down 20 times. The cell lysates were analyzed for protein concentration using the BCA method. 5.5 µg of protein from each cell lysate was loaded onto a protein SDS-PAGE gel. Protein separation by gel-electrophoresis, and subsequent western blot analysis, were performed as described above. The primary antibody incubation lasted overnight with anti-β-actin (Cell Signaling, Danvers MA, Ref: 4970S, Lot: 18) and anti-FLAG-Tag (Cell Signaling, Danvers MA, Ref: 14793S, Lot: 5) primary antibodies. Each antibody was diluted 1:500.

### Identification of METTL7A and METTL7B in human liver microsomes

Pooled 50-donor human liver microsomes (Fisher Scientific, Hampton NH, Ref: HMMCPL, Lot: ADD) and pooled 50-donor human liver cytosol (Fisher Scientific, Hampton NH, Ref: HMMCPL, Lot: PL028-J) were protein normalized, and 40 µg of protein from each subcellular fraction was loaded onto a NuPAGE™ 4 to 12%, Bis-Tris protein gel. Protein separation by gel-electrophoresis, and subsequent western blot analysis were performed as described above. Two western blots were developed, one stained with an anti-METTL7B (Invitrogen, Carlsbad, CA, Ref: PA5-58478, Lot: XC3518324A) and the other with anti-METTL7A primary antibody (Origene, Rockville MD, Ref: TA346478, Lot: Qa2850). These antibodies were diluted 1:1000 and 3:1000, respectively.

### Analysis of DCMB inhibition of TMT activity in liver cellular fractions

50-donor pooled HLM and human liver cytosol to 1 mg/mL were protein normalized. captopril was added to a final concentration of 5 mM, DCMB was added to a final concentration of 100 µM, and SAM was added to a final concentration of 100 µM to initiate the reaction. The reaction was incubated for 45 min at 37 °C. Following incubations, all samples were processed and then analyzed by LC-MS/MS, using the same sample prep and LC-MS/MS method described above.

### Quantification of METTL7A and METTL7B protein levels in individual donor human liver microsomes

Microsomes were prepared by thawing individual donor tissue samples on ice in homogenization buffer (50 mM KPi pH 7.4, 250 mM sucrose, 1 mM EDTA, and 1X Halt protease inhibitor cocktail). The thawed tissue was transferred to a sterile petri dish on ice and a razor blade was used to remove fibrotic tissue. Next, the tissues samples were placed in a 55 mL Dounce homogenizer and homogenized using 4-6 passes with a powered Teflon pestle. The homogenate was transferred to a centrifuge tube and centrifuged at 8,000 × g for 30 min at 4 °C to pellet all intact cells, nuclei, and mitochondria. The supernatant was then transferred to a fresh tube and centrifuged it at 100,000 × g for 60 minutes at 4 °C. The supernatant containing cytosolic fraction was removed and the pellet containing the microsomal fraction was resuspended in 2 mL of homogenization buffer via Dounce homogenization and then aliquoted and stored at −80 °C until future use.

For the quantitative proteomic analysis of METTL7A and METTL7B protein levels in individual donor HLMs, the protein samples were normalized to 2 mg/mL with homogenization buffer. 30 µL of 2 mg/mL HLM were combined with an equal volume of ammonium bicarbonate 100 mM and 10 µL of DTT (250 mM). The samples were then incubated at 37 °C for 30 min. The samples were allowed to cool back to room temperature followed by the addition of 10 µL of iodoacetamide (IAA: 500 mM iodoacetamide in 50 mM ammonium bicarbonate) to each sample. Each sample was then vortexed and incubated for 30 min at room temperature in the dark. 1 mL of crash solution (methanol:chloroform:water, 5:1:4) was added to the samples and centrifuged for 5 min at 16,000 × g and 4 °C. A protein pellet appeared as a disc suspended between the immiscible methanol/water and chloroform layers. Both layers were carefully aspirated and then the protein was allowed to air dry for 10 min. Methanol (0.5 mL) was added to the pellet. Samples were centrifuged for 5 min at 8,000 × g and 4 °C, the methanol was aspirated, and the pellet was allowed to air dry for 30 min followed by the addition of 20 µL of trypsin, dissolved in 50mM ammonium bicarbonate at 1:80 trypsin:protein ratio (w/w). The protein pellet was digested by the trypsin for 16 h while incubating at 37 °C and shaking at 300 rpm. After digestion, 20 µL of LC/MS buffer (80% acetonitrile, with 0.5% formic acid that contains the relative heavy peptides) was added to each sample. The samples where then centrifuged for 10 min at 4,000 x g and 4 °C. the supernatant was transferred and analyzed by LC-MS/MS.

A UPLC-MS/MS (SCIEX Triple Quadrupole 6500 system (Framingham, WA) coupled to an ACQUITY UPLC system (Waters Technologies, Milford, MA) was utilized for proteomic analysis and 5 µl of each sample was injected onto the column (ACQUITY UPLC CSH 1.7 μm, C18; 100×2.1 mm, Waters, Milford, MA). A gradient method (0.3 mL/min) with mobile phase A consisting of 0.1% formic acid in water and mobile phase B consisting of 0.1% formic acid in acetonitrile was used for peptide separation.

Two surrogate peptides purchased from Thermo Fisher Scientific (Waltham, MA) were chosen separately for METTL7A and METTL7B. The relevant transitions for each peptide are shown in supplementary table 01. Analyst 1.6 software (Sciex, USA) was used for peak integration and data analysis. Peakview 2.0 was applied for checking the quality of peaks and initial method development. Chromatographic integration and peak area analysis were performed by using Skyline 20.0.1.31 (University of Washington, Seattle, WA).

### Determination of TMT activity across HLMs from individual donors

Individual liver donor microsomes diluted to 0.25 mg/mL in KPi reaction buffer (50 mM KPi pH 7.0, 20% glycerol, 150 mM NaCl, 10 mM CHAPS) were incubated with 250 µL TSL and 100 µL SAM for 30 min at 37 °C. For the small cohort of individual donor HLMs that were incubated +/-DCMB, the final TSL, DCMB, and SAM concentrations were all 100 µM. The reaction quench and downstream determination of TSL methylation was performed by LC-MS/MS as described above.

### Data analysis

Unless noted otherwise, all experiments were conducted with biological triplicates and all data is reported as the mean ± standard deviation. Statistical significance was determined by a two-tailed unpaired student t-test with a threshold P value of 0.05. Kinetic parameter Km (for TSL and SAM), Ki (for SAH), and IC50 (for DCMB) values were calculated using GraphPad Prism, version 9.4.1 for Windows (GraphPad Software, La Jolla, CA). A two-tailed Spearman’s non-parametric correlation was computed to assess the relationship between two variables with a threshold P value of 0.05.

## Results

### Sequence Similarity and Structural Homology between METTL7A and METTL7B

Both METTL7A and METTL7B are 244 residues long and have a molecular weight of 28 kDa, similar to prior TMT literature (Weisiger and Jakoby, 1979). METTL7A and METTL7B share 75% sequence homology and 60% sequence identity as determined by BLAST analysis (Altschul et al., 1990) (supplementary figure 01). Homology models of METTL7A and METTL7B, produced by AlphaFold (Jumper et al., 2021; Varadi et al., 2022), indicate high structural similarity (Fig. 1).

**Figure. 1.**
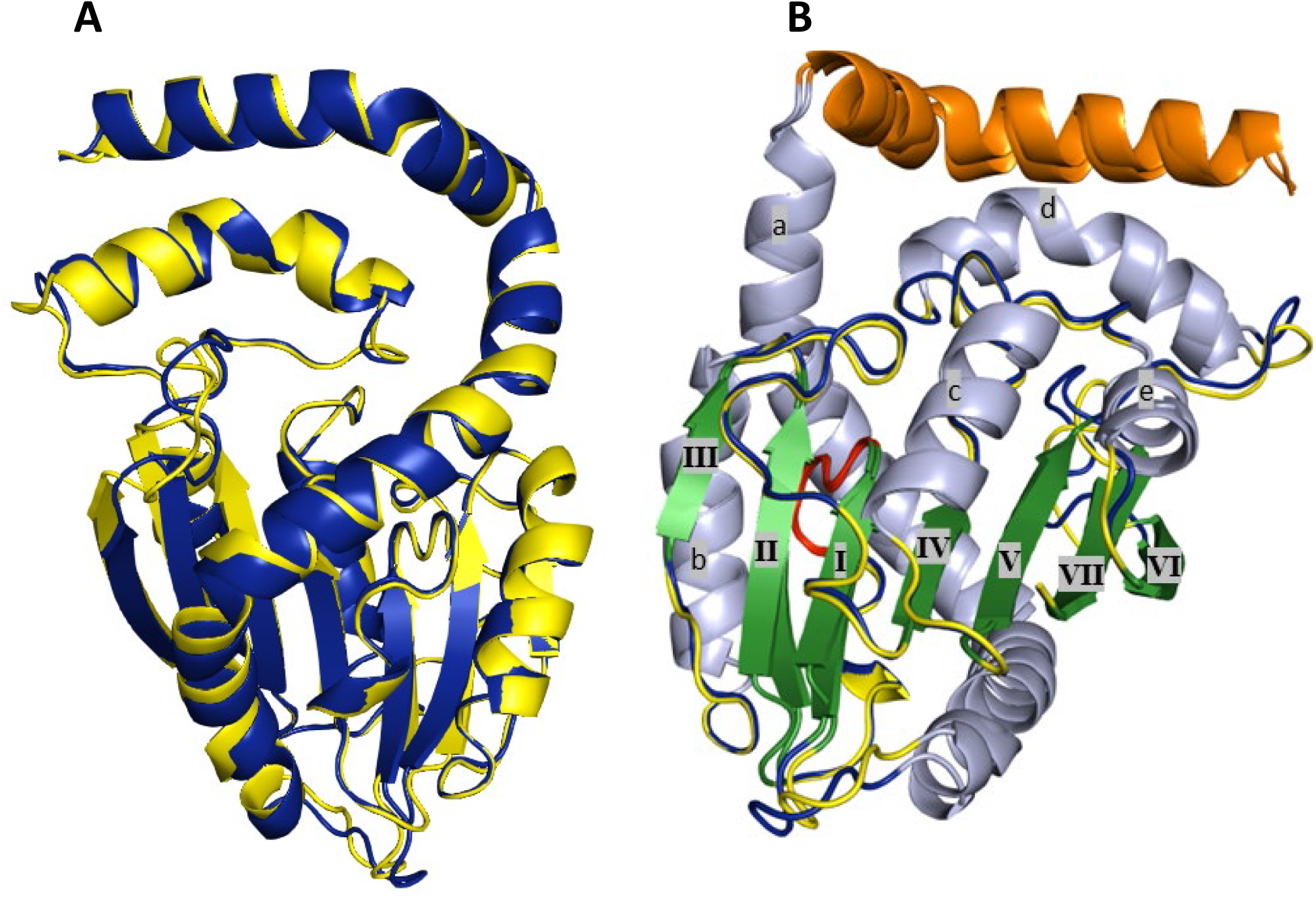
AlphaFold homology models of METTL7A (yellow) and METTL7B (blue) aligned with Pymol alignment function (A). AlphaFold homology models of METTL7A (yellow) and METTL7B (blue) aligned and oriented to best see secondary structural features, with hydrophobic N-terminus colored orange, GXGXG sequence colored red, beta sheets colored green and labeled I-VII by order along the sequence, and alpha helices colored light blue and labeled a-e by order along the sequence (B).

### Expression and purification of His-GST-METTL7A

Consistent with the previous expression of METTL7B (Maldonato et al., 2021), we inserted the METTL7A gene sequence into a pET21 expression plasmid and expressed the protein in *E. coli*. The pET21 expression vector adds a unique tag to the *N*-terminus of the inserted ORF, a 10X histidine-tag (His) followed by a glutathione-*S-*transferase (GST) tag (Fig 2.). The N-terminal GST tag is essential to solubilize METTL7A from the bacterial membrane. The recombinant protein 10XHis-GST-METTL7A (N-GST-METTL7A) has a molecular weight of ∼57.6 kDa. We confirmed the successful purification of N-GST-METTL7A with a visible protein band at ∼58 kDa by SDS-PAGE Coomassie stain (Fig. 2). A band at ∼58 kDa is also visible on a western blot using an anti-METTL7A and anti-GST-antibody; it is not visible using an anti-METTL7B-stained western blot (Fig. 2). The purity of recombinant METTL7A was comparable to the purity we previously achieved for recombinant METTL7B and the lower molecular weight bands that we have previously identified as non-functional GST tag portion of N-GST-METTL7B also co-purified with N-GST-METTL7A. We suspect that the sequence between GST and METTL7A or 7B introduced a feature that leads to the formation of this N-terminal tag portion of this protein construct during the process of bacterial protein expression. There is no concern that the non-specific proteins observed in the purified protein eluent are involved in TMT activity as we have demonstrated previously with a non-functional D98A mutant METTL7B that there is no TMT activity in purification eluent or bacterial lysate without a functional, expressed thiol methyltransferase (Maldonato et al., 2021).

**Fig. 2.**
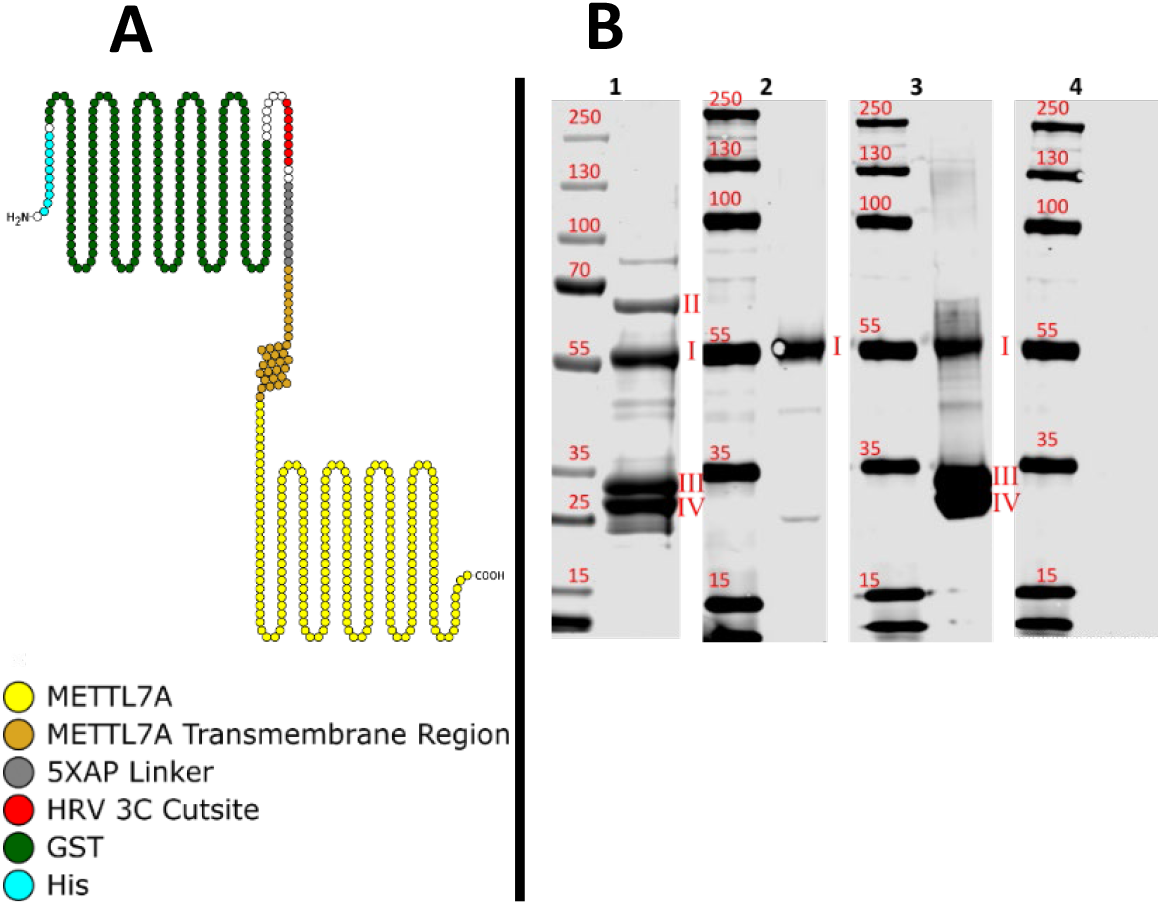
Cartoon representation of N-GST-METTL7A with features highlighted and identified in legend (A) Figure produced with Protter (Omasits et al., 2014). SDS-PAGE Coomassie stain (B. 1), anti-METTL7A western blot (B. 2), anti-GST western blot (B. 3), and anti-METTL7B western blot (B. 4) of purified N-GST-METTL7A. Labeled bands are N-GST-METTL7A (I), a contaminant chaperone protein (II), and GST devoid of METTL7A (III, IV).

### Validation of N-GST-METTL7A *S-*methyltransferase activity

Purified N-GST-METTL7A methylated captopril and TSL (Fig. 3) in a time- and protein-dependent fashion. Like N-GST-METTL7B, purified N-GST-METTL7A required preincubation with dimyristoyl-sn-glycero-3-PG (DMPG)) liposomes for activity (supplementary figure 02).

**Figure. 3.**
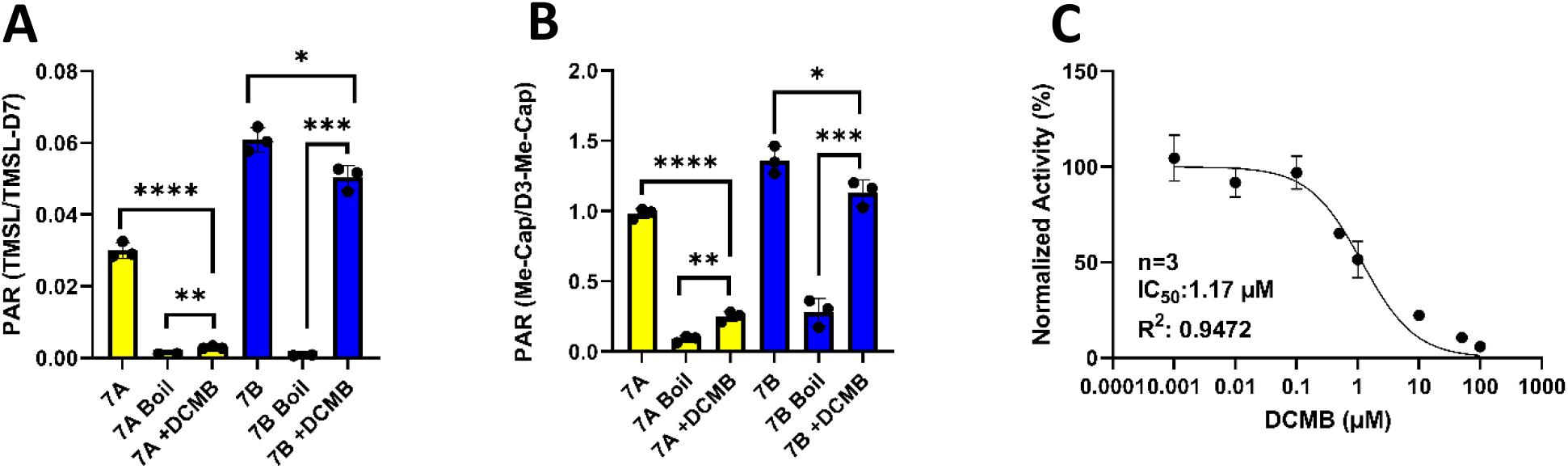
Methyltransferase activity of purified recombinant METTL7A and METTL7B +/-DCMB. Monitoring for methylation of TSL (A) or captopril (B) confirms that DCMB inhibits METTL7A but not METTL7B. An IC50 for the inhibition of purified recombinant METTL7A by DCMB was determined by single time-point activity assays with increasing concentrations of DCMB (C). Data are reported in triplicate (mean +/-S.D.). Significance was determined using an unpaired two-tailed t-test comparing bracketed treatments ****P < 0.0001. ***P < 0.001. **P < 0.01. The IC50 curve enzyme activity assay reactions were conducted with TSL just above Km and the IC50 curve was fitted using GraphPad prism’s equation for [Inhibitor] vs. normalized response.

### Inhibition of METTL7A by DCMB

DCMB potently inhibits N-GST-METTL7A but not N-GST-METTL7B (Fig. 3). An IC50 of 1.2 µM was determined for DCMB and METTL7A, using TSL as a reporter substrate (Fig. 3). There is virtually no 7α-thiomethylspironolactone (TMSL) observed in the inactivated boil controls (supplementary figure 03), and because of this, we concluded TSL to be a suitable probe substrate for further METTL7 characterization experiments.

### Competitive inhibition of N-GST-METTL7A by *S-*adenosyl-homocysteine

We determined the Km value for the methyl donating cofactor SAM by measuring the formation of TMSL by LC-MS/MS (Fig. 4). The SAM activity curve followed Michaelis-Menten kinetics. The Km for SAM was 53.73 µM. Analysis of the METTL7A homology model generated by AlphaFold indicated that this protein is a class I small molecule methyltransferase (supplementary figure 04). Class I methyltransferases have a conserved binding site with a high affinity for SAM and the demethylated cofactor *S-*Adenosyl-homocysteine (SAH) (Schubert et al., 2003). As expected with a class I small molecule methyltransferase, SAH inhibited TSL methylation in a concentration-dependent manner and exhibited a competitive mechanism of inhibition with respect to SAM (Fig. 4), with a KI of 53.9 µM. We measured all kinetic parameters under linear conditions for incubation time and protein concentration (supplementary figure 05). For SAM and SAH kinetic analysis, a saturating amount of TSL was added for these assays to be under conditions of no substrate depletion.

**Figure. 4.**
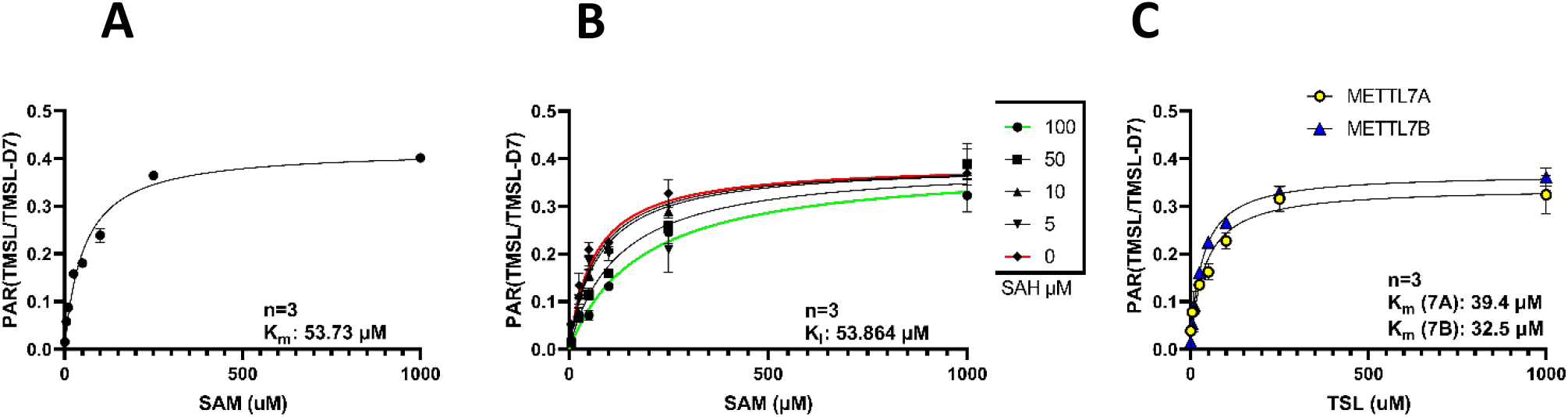
TSL methylation by N-GST-METTL7A at various concentrations of SAM without (A) and with various concentrations of SAH (B); and single time-point kinetics profiles of N-GST-METTL7A or N-GST-METTL7B at various concentrations of TSL (C). Data are reported in triplicate (mean +/-S.D.). TSL and SAM both exhibit saturable Michaelis-Menten kinetics. Kinetic parameter Km and Ki (for SAH) values were obtained by non-linear regression analysis using GraphPad Prism.

### Kinetic analysis of *N*-GST-METTL7A and *N*-GST METTL7B *S*-methylation of TSL

We determined the Km value for TSL by measuring the formation of TMSL for both N-GST-METTL7A and N-GST-METTL7B by LC-MS/MS (Fig. 4). The TSL activity curves for each enzyme followed Michaelis-Menten kinetics. The Km values for TSL methylation by METTL7A and METTL7B were 39.4 and 32.5 µM, respectively. The Km for TSL methylation by recombinant METTL7B by LC-MS/MS, an orthogonal approach to our previous analysis, was similar to what we have previously reported for METTL7B (Maldonato et al., 2021).

### Substrate specificity of N-GST-METTL7A

Substrates for class I small molecule methyltransferases were screened for N-GST-METTL7A methylation with the Promega MTaseGlo kit. In previous screens with this assay kit, METTL7B activity was normalized to dopamine, a compound that showed no activity. Normalization was necessary because METTL7B produced a high response in absence of a substrate compared to a boiled enzyme control. METTL7A, incubated without a substrate, also demonstrated a high response compared to the boiled enzyme. Adding DCMB, as a quench step, at the end of incubation significantly reduced the baseline signal (supplementary figure 06). This observation indicates that METTL7A and METTL7B catalyze the methylation of a compound in the reagents supplied with the kit. As such, it became standard practice to include DCMB in the quench step to prevent further METTL7A activity during downstream signal development steps.

Potential TMT substrates were screened at concentrations at least three times higher than their literature reported Km values to ensure the detection of methylation activity (Weisiger and Jakoby, 1979; Rivett and Roth, 1982; Drummer et al., 1983; Hiemke and Ghraf, 1983; Keith et al., 1984; Keith et al., 1985; Grunewald et al., 1989). METTL7A preferentially methylated exogenous thiol-containing compounds and did not methylate the endogenous thiols tested; cysteine and glutathione (Fig. 5). METTL7A did not methylate compounds that are known O- and N-methyltransferase substrates. Mertansine, a thiol-containing microtubule inhibitor, was significantly methylated above baseline. We also tested *S-*methyl mertansine as a negative control, and the response was not significantly above baseline. We assume that mertansine can only be methylated on its thiol moiety. Because we assume that *S-*methyl mertansine has no site for further methylation, we used *S*-methyl mertansine activity as a non-substrate baseline control for screening other potential substrates with the Promega MTaseGlo assay.

**Figure. 5.**
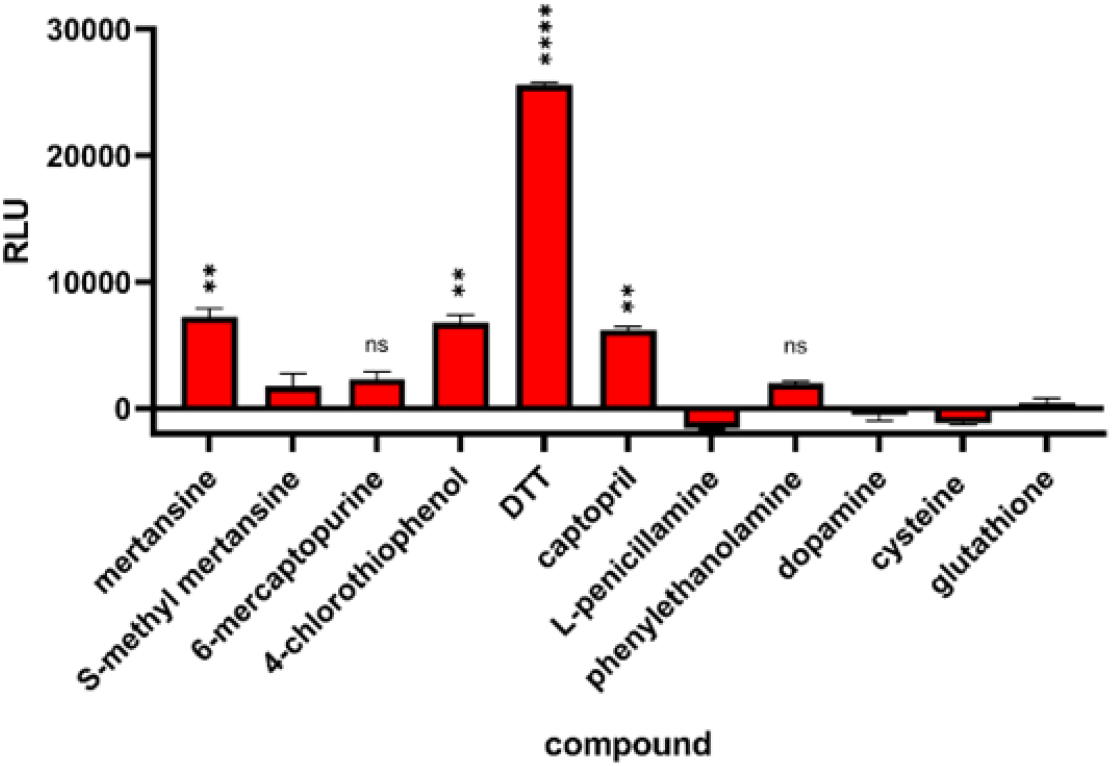
Semi-quantitative screening of select small molecule methyltransferase probe substrates. Multiple methyltransferase probe substrates were incubated at saturating concentrations for 1 h with N-GST-METTL7A. Formation of SAH (indication of methyl transfer) was converted/directly linked to luminescence and measured via the Promega MTaseGlo kit. Enzyme activity was determined after normalizing to compound-specific and general assay background signal. All data are presented as the mean ± s.d. Significance was determined using unpaired two-tailed t test comparing the baseline subtracted response of each compound that reports above zero with the response of S-methyl mertansine. ****P < 0.0001. ***P < 0.001. **P < 0.01.

### Modulating METTL7A and METTL7B gene expression in HepG2 cells and its effect on thiol methylation activity

HepG2 cells were treated with scrambled negative control siRNA, METTL7A-specific small interfering RNA (siRNA), METTL7B-specific siRNA, or a combination of both. METTL7A-specific siRNA exclusively reduced METTL7A gene expression by 90%, whereas METTL7B-specific siRNA reduced the expression of METTL7B and METTL7A by 90%and 30%, respectively. The non-specific reduction in METTL7A gene expression using a METTL7B-specific siRNA is likely caused by siRNA design. The METTL7B specific siRNA shares 60% sequence complementary with a METTL7A coding mRNA segment, potentially inducing modest suppression (supplementary figure 07). The combined treatment of METTL7A- and METTL7B-specific siRNA reduced the expression of both genes by 90% (Fig. 6). After incubation with siRNA for 48 h, cell media was replaced with media containing either TSL or captopril. The methylated metabolites for each compound were measured in the culture media by LC-MS/MS. We observed the largest decrease in TSL and captopril methylation in the combined siRNA-treated cell media compared to cells treated with control siRNA. The smallest decrease was observed in METTL7B-specific siRNA-treated cell media (Fig. 6).

**Figure. 6.**
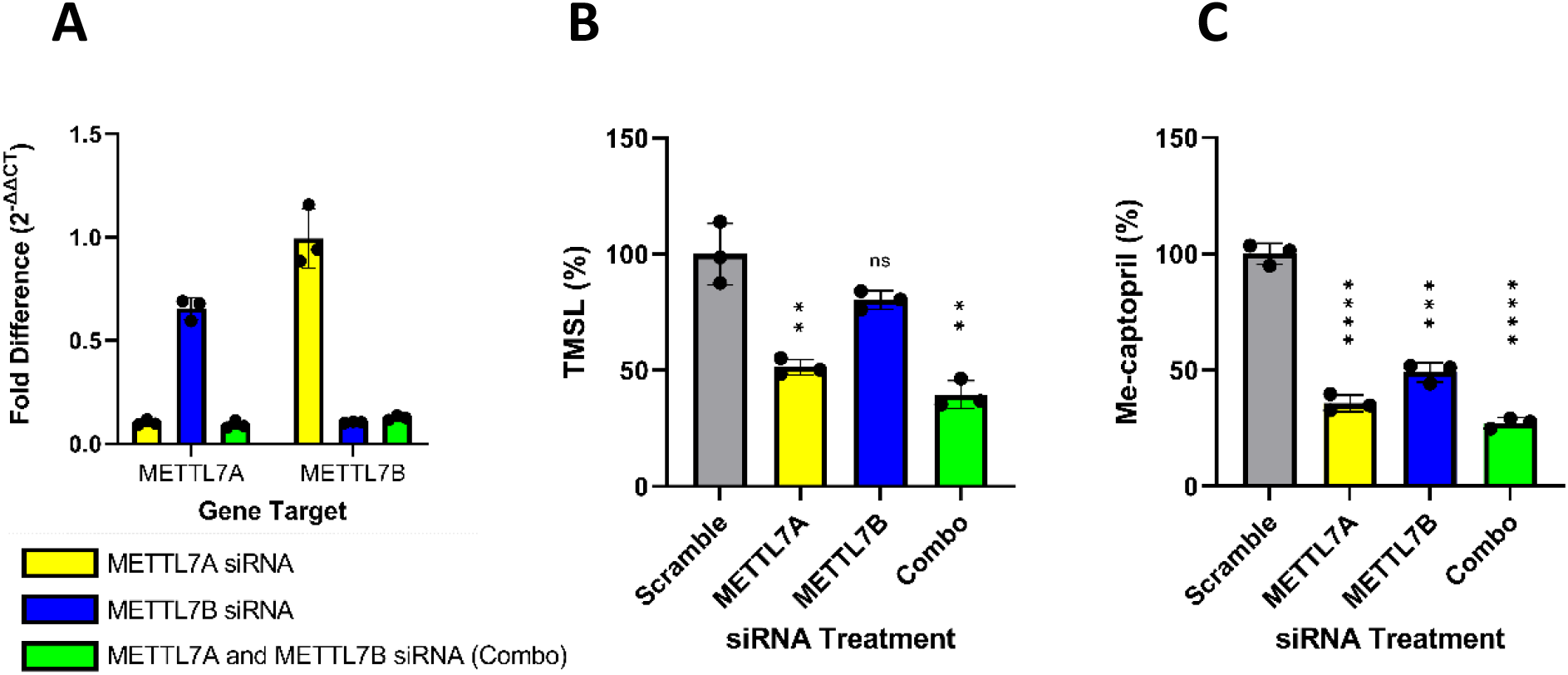
Knockdown of *METTL7A* and *METTL7B* in HepG2 cells. RT-PCR data measuring fold difference of *METTL7A* or *METTL7B* expression following 48-hour siRNA treatment with *METTL7A-* and/or *METTL7B*-targeting siRNA (A). Percent TMSL (B) and S-methyl captopril (C) formed in cell media of siRNA-treated HepG2 cells. All data are presented as the mean ±s.d. Significance was determined using an unpaired two-tailed t-test. ****P < 0.0001. ***P < 0.001. **P < 0.01.

### Modulating METTL7A and METTL7B protein levels in HeLa cells and its effect on thiol methylation activity

HeLa cells were incubated with pCMV expression vectors encoding FLAG-tagged METTL7A, FLAG-tagged METTL7B, or an empty insert to act as a negative control. Protein expression of METTL7A or METTL7B was determined in cell lysate by protein normalized anti-FLAG western blotting. An anti-β-actin antibody was included as a co-stain for loading control. FLAG-tagged METTL7A and METTL7B are visible at the correct molecular weights, only in lysates of cells incubated with the appropriate expression vector (Fig. 7). No FLAG-tagged protein is visible in the empty control vector-treated cells. Cells expressing FLAG-tagged METTL7A or FLAG-tagged METTL7B had significantly increased TSL methylation compared with empty vector control-treated cells, determined by LC-MS/MS (Fig. 7). With the addition of DCMB, only METTL7A overexpressing cells showed a significant reduction in TSL methylation compared to cells not treated with DCMB.

**Figure. 7.**
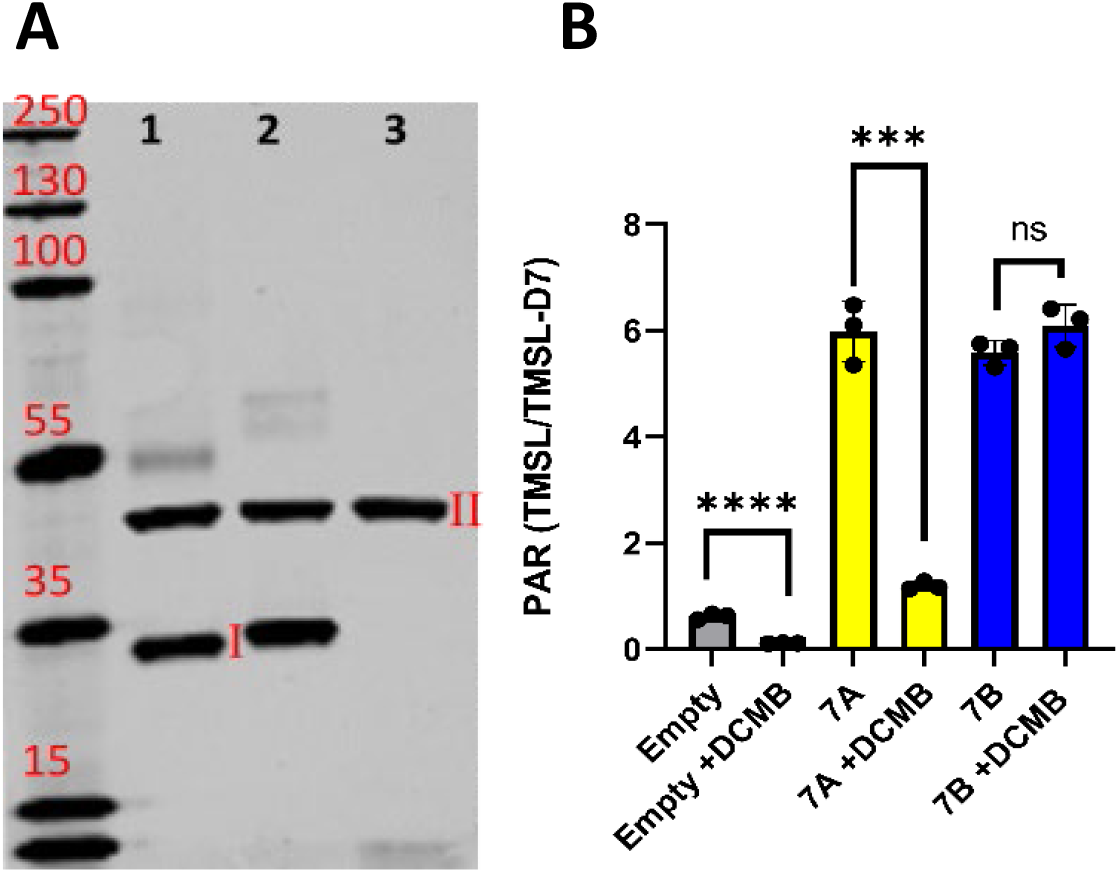
Overexpression of METTL7A and METTL7B in HeLa cells. Anti-flag and anti-β-actin western blot verify the overexpression of flag-tagged proteins in METTL7A (A. 1) and METTL7B (A. 2) which show a band at I, compared to cells treated with empty vector control (A 3.) which do not. Labeled bands are flag-tagged METTL7A and B (I), and β-actin (II). Formation of TMSL in HeLa cells overexpressing METTL7A or METTL7B following a 24-hour incubation with TSL +/-DCMB confirms that only METTL7A is inhibited by DCMB (B). All data are presented as the mean ±s.d. Significance was determined using unpaired two-tailed t-test. ****P < 0.0001. ***P < 0.001. **P < 0.01.

### Subcellular localization of METTL7A and METTL7B in human liver

Using 50-donor pooled HLMs, we observed bands at the molecular weight of 28 kDa in anti-METTL7A and anti-METTL7B stained western blots (Fig. 8). We observed similar bands in the pooled liver cytosolic fraction, albeit at much lower intensities.

**Figure. 8.**
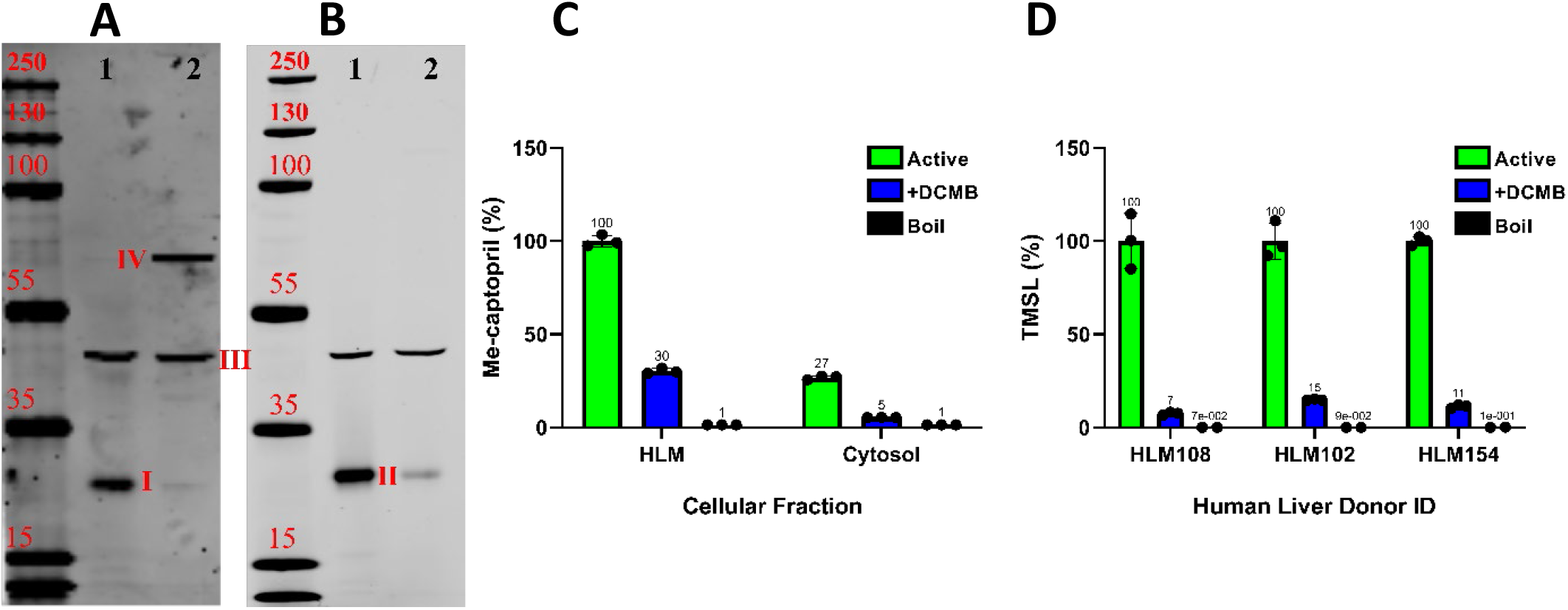
Anti-METTL7A (A) and anti-METTL7B (B) western blots of human liver microsomes (1) and cytosol (2). Labeled bands are METTL7A (I), METTL7B (II), and β-actin (III), the band at IV is a mystery band that shows up by anti-METTL7A antibody in the cytosolic liver subcellular fraction. Relative formation of S-methyl-captopril in human liver microsomes, cytosol, and S9 fractions +/-DCMB (C) confirms literature findings that the majority of TMT activity is associated with microsomes. Finally, individual donors incubated with TSL and SAM +/-DCMB (D) confirms literature findings that the majority of TMT activity is lost with coincubation of microsomal TMT activity assay with DCMB. All treatments performed in triplicate except for figure D boil controls which were performed in duplicate. All data are presented as the mean ±s.d.

We determined protein-normalized TMT activity in the 50-donor pooled microsomal or cytosolic fractions by measuring captopril methylation via LC-MS/MS analysis (Fig. 8). HLMs contained the highest TMT activity, while lowest TMT activity was observed in liver cytosol, which had 30% TMT activity compared to HLMs.

We determined the effect of DCMB on TMT activity in pooled HLMs and cytosol by adding saturating concentrations of DCMB during captopril methylation assays (Fig. 8). There was ≥70% reduction in TMT activity in HLMs and liver cytosol when incubated with DCMB. In a similar inhibition experiment, we screened individual-donor HLMs for TMT activity +/-DCMB. However, instead of using captopril, we used TSL as the probe substrate (Fig. 8). Saturating concentrations of DCMB, added during TSL methylation assays, reduced each individual-donor HLM’s TMT activity by >80%.

### Quantifying of METTL7A and METTL7B protein levels in HLMs and correlation with *S*-methylation activity

We quantified METTL7A and METTL7B protein levels in 19 adult single-donor HLMs by LC-MS/MS quantitative proteomics (Fig. 9). METTL7A protein levels are, on average, 3-fold higher than METTL7B levels. We also determined TSL methylation activity in these same donor HLMs (Fig. 9). There was an 8-fold difference between the highest and lowest activity among the donors.

**Figure. 9.**
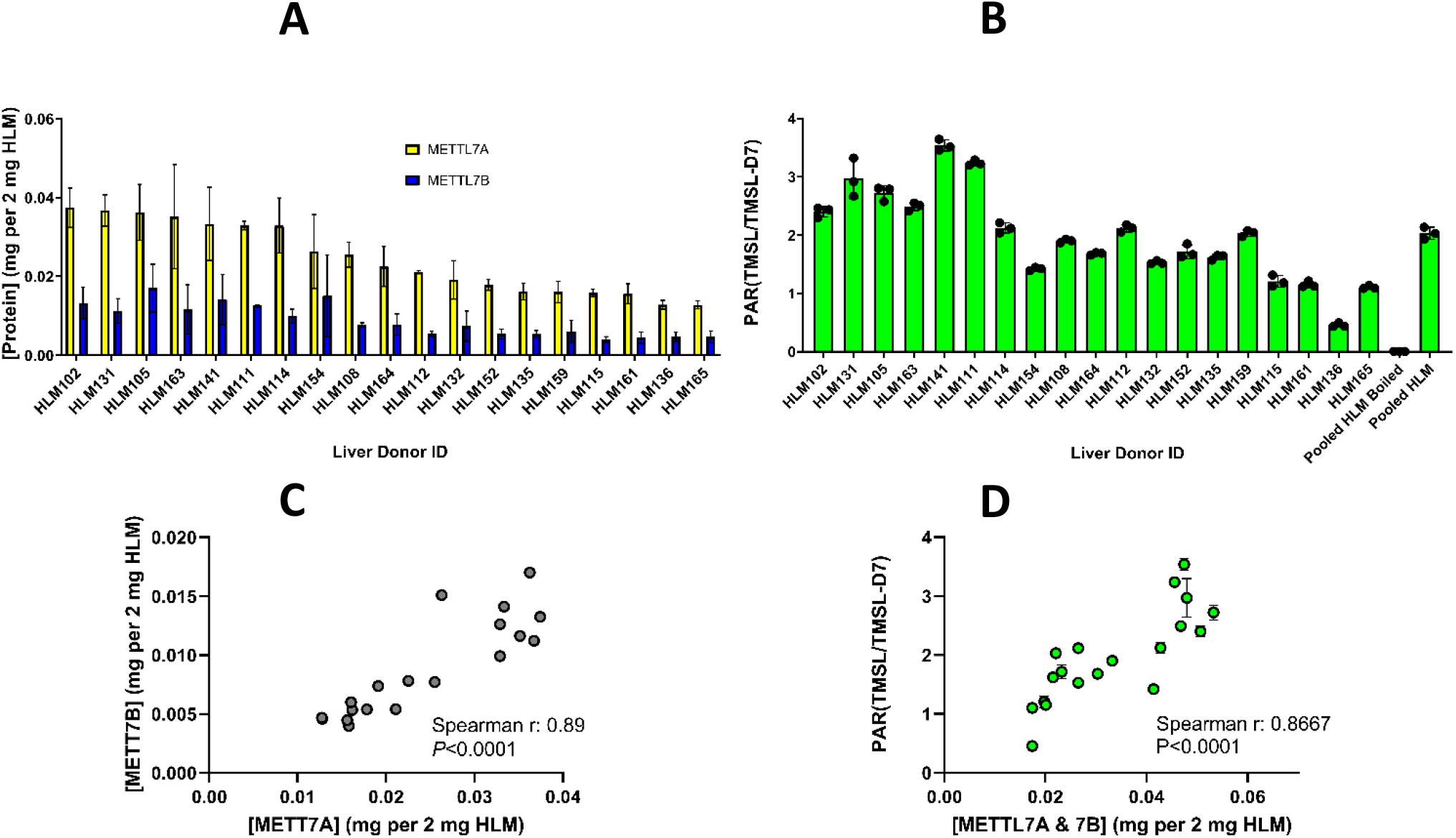
METTL7A and METTL7B protein expression is correlated to each other and overall TMT activity (A). Protein levels of METTL7A and METTL7B are reported normalized to total protein per digestion. Methyltransferase activity of single-donor HLMs incubated with TSL, and SAM (B) compared with commercially available 50-donor pooled HLMs. METTL7A and METTL7B protein levels are strongly correlated (C). TMT activity is correlated with total METTL7A and METTL7B protein levels (D). Data are presented as the mean ±s.d. Nonparametric Spearman correlation was determined using GraphPad Prism.

We computed a non-parametric Spearman’s correlation to assess the relationship between METTL7A and METTL7B protein levels. The Spearman correlation coefficient was 0.85, and the P-value was < 0.0001 (Fig. 9). This indicates a positive correlation between METTL7A and METTL7B. We also calculated a Spearman correlation coefficient to assess the relationship between relative TSL methylation activity and the combined protein levels of METTL7A and METTL7B. The Spearman correlation coefficient for this analysis was 0.8667, and the P-value was < 0.0001 (Fig. 9). This indicates a positive correlation between METTL7A and METTL7B protein levels and relative TMT activity across individual donor adult HLMs.

## Discussion

The key finding in this investigation is that METTL7A is a thiol methyltransferase and, along with METTL7B, accounts for most of *S*-thiol methylation activity in HLMs. We will refer to METTL7A and METTL7B as TMT1A1 and TMT1B1, respectively, for the rest of the discussion and use *METTL7A* and *METTL7B* when referring to the genes. TMT1A1 can selectively methylate exogenous alkyl and phenolic thiols, and is inhibited by DCMB, the historic TMT inhibitor with an IC50 of 1.17 µM which is similar to the IC50 of 0.45 µM for TMT determined in HLMs (Glauser et al., 1993). Therefore, although TMT1B1 is a thiol methyltransferase, it cannot entirely be responsible for TMT activity because DCMB has no effect on its activity. TMT1A1 shares 60% sequence identity with TMT1B1 and has a similar substrate specificity, yet only TMT1A1 is potently inhibited by DCMB. Further work is required to determine the structural features in TMT1A1 that confer DCMB selectivity compared to TMT1B1.

Recombinant TMT1A1 transferred a methyl group to several exogenous thiol-containing compounds, including DTT, mertansine, captopril, and 6-chlorothiophenol in a general methyltransferase substrate screening assay. Similar to prior reports for TMT, TMT1A1 did not methylate endogenous thiols including cysteine and glutathione or other small molecules specific for N- and O-methyl transferases (Weisiger and Jakoby, 1979).

TMT1A1 behaves like a class I small molecule methyltransferase determined by a series of kinetic experiments. TMT1A1 utilizes SAM as the methyl donor and methylates TSL following classic Michaelis-Menten kinetics. It is competitively inhibited by SAH, as evidenced by higher SAM concentrations overcoming SAH inhibition. The Michaelis-Menten constant (Km) for SAM and TSL are 53.73 µM and 39.4 µM, respectively; in close agreement with the values of 43 µM and 27 µM previously reported in the literature (Weinshilboum et al., 1979; Keith et al., 1984). In addition, we compared the kinetic curves of TSL methylation by TMT1A1 and TMT1B1 and established that both recombinant enzymes have similar Km values for TSL. The Km value for TMT1B1 mediated methylation of TSL, reported by LC-MS/MS, was 32.5 µM, close to the value determined in our previous study, which instead utilized the Promega MTase-Glo™ Methyltransferase Assay to report on the methylation of TSL.

*METTL7A* and *METTL7B* gene knockdown experiments in HepG2 cells caused a significant decrease in *S*-methylation activity, while overexpression of *METTL7A* and *METTL7B* in HeLa cells caused a significant increase in TMT activity, as expected. Only HeLa cells overexpressing TMT1A1 had reduced TMT activity in the presence of DCMB confirming that the specificity towards TMT1A1 is not due to a difference in the in vitro expression or incubation systems.

TMT activity is described in the literature as most abundant in liver microsomes (Weisiger and Jakoby, 1980; Weisiger et al., 1980; Drummer et al., 1983), and an RNA sequencing analysis reported that like TMT, *METTL7A*, and *METTL7B* expression levels are highest in the human liver (Karlsson et al., 2021) compared to other tissues. By western blot analysis, we determined that TMT1A1 and TMT1B1 are present in the human liver and are associated with HLM. We also confirmed that TMT activity is highest in HLM and lowest in the liver cytosol consistent with previous reports (Bremer and Greenberg, 1961; Borchardt and Cheng, 1978; Weisiger and Jakoby, 1979; Pacifici et al., 1991).

Saturating concentrations of DCMB added to a TMT activity assay performed in HLMs, results in ≥70% decrease in TMT activity towards captopril methylation and >80% decrease in activity towards TSL methylation. Because TMT1A1 is potently inhibited by DCMB while TMT1B1 is not, we believe that we can attribute most of the TMT activity in HLMs to TMT1A1, and the activity remaining with saturating concentrations of DCMB can be attributed to TMT1B1. Finally, we measured the protein levels of TMT1A1 and TMT1B1 by quantitative proteomics. TMT1A1 protein levels are, on average, 3-fold greater than TMT1B1 but the two proteins were highly correlated. TMT activity was determined for each donor and correlated well with the sum of TMT1A1 and TMT1B1 protein levels, as seen in figure 9.

The high TMT1A1 and TMT1B1 protein levels in the liver compared to other tissue and their association with the microsomal cellular fraction parallel other drug-metabolizing enzymes like cytochrome P450 enzymes (Šrejber et al., 2018). Other class I small molecule methyltransferases such as catechol-O-methyltransferase (COMT), phenylethanolamine *<u>N-</u>*methyltransferase (PNMT), and nicotinamide N-methyltransferase (NNMT) have very limited examples of exogenous substrates but clear examples of endogenous ones—dopamine, noradrenaline, and nicotinamide, respectively (Aksoy et al., 1994; Drinkwater et al., 2009; Júlio-Costa et al., 2013). In this regard, it is reasonable to consider TMT1A1 and TMT1B1 as drug metabolizing enzymes, but they could also have an endogenous role, such as the methylation of hydrogen sulfide or other endogenous biomolecules. We have previously reported that TMT1B1 methylates hydrogen sulfide (Maldonato et al., 2021), and preliminary data from our lab suggests that TMT1A1 also methylates hydrogen sulfide (data not shown).

While the endogenous function of TMT1A1 and TMT1B1 remains unknown, their dysregulation is continually reported in different diseases, especially cancer. In one large-cohort multi-cancer, multiomics study, higher *METTL7A* expression was determined to have the most favorable survival across cancers compared to other METTLs (Campeanu et al., 2021). *METTL7B*, antithetically, is significantly upregulated across most cancers (Jiang et al., 2021). TMT1B1 regulates the epithelial-mesenchymal transition of thyroid cancer cells in the presence of TGF-β, an important regulator of cancer immunosuppression (Ye et al., 2019; Chen et al., 2021; Xiong et al., 2021); and in multiple cell lines, TMT1B1 promotes G1 to S phase transition in cancer cells (Liu et al., 2020; Li et al., 2021). *METTL7B* overexpression is attributed to proliferation and tumorigenesis, which suggests an oncogenic role in cancer progression. Conversely, the consistent lower expression of *METTL7A* across cancers would suggest a tumor-suppressing role. Although the nature of TMT1A1 and TMT1B1’s involvement in cancers is not well understood, the high level of dysregulation of *METTL7A* and *METTL7B* might impact the metabolism of thiol-containing chemotherapeutics by cancer cells.

Several papers have postulated that TMT1A1 and TMT1B1 are capable of RNA methylation (Wang et al., 2022), which is an interesting endogenous function and worth exploring in future work; however, microsomal association of these enzymes is more indicative of a role in drug metabolism and the metabolism of other thiol-containing biomolecules.

Our findings are consistent with the hypothesis that TMT1A1 and TMT1B1, together, are responsible for most of the microsomal TMT activity. Future work will focus on identifying endogenous thiol-containing substrates for TMT1A1 and TMT1B1, examining their role in tumorigenesis and determining the structural features that account for DCMB selective inhibition of TMT1A1.

## Acknowledgments

The authors wish to thank Scott Edgar for his advice and contribution to mass spec assay development.

We would also like to thank Ian Levasseur for his help with protein purification.

## Authorship Contributions

*Participated in research design:* Russell, Maldonato, Shi, Chau, Totah.

*Conducted experiments:* Russell, Chau, Shi, Maldonato

*Contributed new reagents or analytic tools:* Russell, Shi, Chau.

*Performed data analysis:* Russell, Chau, Shi, Totah.

*Wrote or contributed to the writing of the manuscript:* Russell, Chau, Shi, Maldonato, Totah.

