## Supplementary File for "METTL7A and METTL7B are responsible for *S*-thiol methyl transferase activity in liver"

| Peptide list | parent ions | fragmentations |
|---|---|---|
| sp\|Q9H8H3\|MET7A_HUMAN.VTC[CAM]IDPNPNFEK.+2y9.light | 717.34 | 1073.53 |
| sp\|Q9H8H3\|MET7A_HUMAN.VTC[CAM]IDPNPNFEK.+2y8.light | 717.34 | 960.44 |
| sp\|Q9H8H3\|MET7A_HUMAN.VTC[CAM]IDPNPNFEK.+2y7.light | 717.34 | 845.42 |
| sp\|Q9H8H3\|MET7A_HUMAN.VTC[CAM]IDPNPNFEK.+2y5.light | 717.34 | 634.32 |
| sp\|Q9H8H3\|MET7A_HUMAN.VTC[CAM]IDPNPNFEK.+2y9.heavy | 721.35 | 1081.54 |
| sp\|Q9H8H3\|MET7A_HUMAN.VTC[CAM]IDPNPNFEK.+2y8.heavy | 721.35 | 968.46 |
| sp\|Q9H8H3\|MET7A_HUMAN.VTC[CAM]IDPNPNFEK.+2y7.heavy | 721.35 | 853.43 |
| sp\|Q9H8H3\|MET7A_HUMAN.VTC[CAM]IDPNPNFEK.+2y5.heavy | 721.35 | 642.33 |
| MET7A.FYPPGC[CAM]R.+2y5.light | 448.71 | 586.28 |
| MET7A.FYPPGC[CAM]R.+2y4.light | 448.71 | 489.22 |
| MET7A.FYPPGC[CAM]R.+2y5+2.light | 448.71 | 293.64 |
| MET7A.FYPPGC[CAM]R.+2y4+2.light | 448.71 | 245.12 |
| MET7A.FYPPGC[CAM]R.+2y5.heavy | 453.71 | 596.28 |
| MET7A.FYPPGC[CAM]R.+2y4.heavy | 453.71 | 499.23 |
| MET7A.FYPPGC[CAM]R.+2y5+2.heavy | 453.71 | 298.65 |
| MET7A.FYPPGC[CAM]R.+2y4+2.heavy | 453.71 | 250.12 |
| sp\|Q6UX53\|MET7B_HUMAN.VTC[CAM]LDPNPHFEK.+3y7.light | 486.23 | 868.43 |
| sp\|Q6UX53\|MET7B_HUMAN.VTC[CAM]LDPNPHFEK.+3y11+2.light | 486.23 | 679.31 |
| sp\|Q6UX53\|MET7B_HUMAN.VTC[CAM]LDPNPHFEK.+3y10+2.light | 486.23 | 628.79 |
| sp\|Q6UX53\|MET7B_HUMAN.VTC[CAM]LDPNPHFEK.+3y7+2.light | 486.23 | 434.72 |
| sp\|Q6UX53\|MET7B_HUMAN.VTC[CAM]LDPNPHFEK.+3y7.heavy | 488.91 | 876.45 |
| sp\|Q6UX53\|MET7B_HUMAN.VTC[CAM]LDPNPHFEK.+3y11+2.heavy | 488.91 | 683.32 |
| sp\|Q6UX53\|MET7B_HUMAN.VTC[CAM]LDPNPHFEK.+3y10+2.heavy | 488.91 | 632.8 |
| sp\|Q6UX53\|MET7B_HUMAN.VTC[CAM]LDPNPHFEK.+3y7+2.heavy | 488.91 | 438.73 |
| MET7B.ELFSQIK.+2y6.light | 432.74 | 735.44 |
| MET7B.ELFSQIK.+2y5.light | 432.74 | 622.36 |
| MET7B.ELFSQIK.+2y4.light | 432.74 | 475.29 |
| MET7B.ELFSQIK.+2y6+2.light | 432.74 | 368.22 |
| MET7B.ELFSQIK.+2y6.heavy | 436.75 | 743.45 |
| MET7B.ELFSQIK.+2y5.heavy | 436.75 | 630.37 |
| MET7B.ELFSQIK.+2y4.heavy | 436.75 | 483.3 |
| MET7B.ELFSQIK.+2y6+2.heavy | 436.75 | 372.23 |

**Supplementary table 01.** Table showing transitions for heavy labeled and unlabeled surrogate peptides for METTL7A and METTL7B.

```
        #    1         10          20          30          40          50          60
METTL7A      MELTIFILRL AIYILTFPLY LLNFLGLWSW ICKKWFPYFL VRFTVIYNEQ MASKKRELFS
             M++ + +L L ++ +LT PL  L   LG W   +CK  FPY   +  T   N   M SKKRELFS
METTL7B      MDILVPLLQL LVLLLTLPLH LMALLGCWQP LCKSYFPYLM AVLTPKSNRK MESKKRELFS
        #    61        70          80          90          100         110         120
METTL7A      NLQEFAGPSG KLSLLEVGCG TGANFKFYPP GCRVTCIDPN PNFEKFLIKS IAENRHLQFE
              +    G SG K+ LLE+GCG TGANF FYPP GCRVTC+DPN P FEKFL KS  AENRHLQ+E
METTL7B      QIKGLTGASG KVALLELGCG TGANFQFYPP GCRVTCLDPN PHFEKFLTKS MAENRHLQYE
        #    121       130         140         150         160         170         180
METTL7A      RFVVAAGENM HQVADGSVDV VVCTLVLCSV KNQERILREV CRVLRPGGAF YFMEHVAAEC
             RFVVA GE M +Q+ADGS DV VVCTLVLCSV     ++L EV  RVLRPGG+  +F EHVA
METTL7B      RFVVAPGEDM RQLADGSMDV VVCTLVLCSV QSPRKVLQEV RRVLRPGGVL FFWEHVAEPY
        #    181       190         200         210         220         230         240
METTL7A      STWNYFWQQV LDPAWHLLFD GCNLTRESWK ALERASFSKL KLQHIQAPLS WELVRPHIYG
               W + WQQV  +P W+ + D GC LTRE WK  LE A FS +    +   PL  W  V PHI G
METTL7B      GSWAFMWQQV FEPTWKHIGD GCCLTRETWK DLENAQFSEI QMERQPPPLK WLPVGPHIMG
        #       244
METTL7A      YAVK
              AVK
METTL7B      KAVK
```

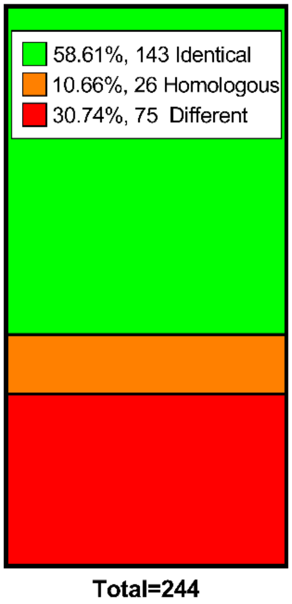


Supplementary Figure **01.** Basic local alignment results for METTL7A compared with METTL7B. Residues 22-244 were returned with alignment results from NCBI’s BLAST search algorithm while residues 1-21 were added by hand.

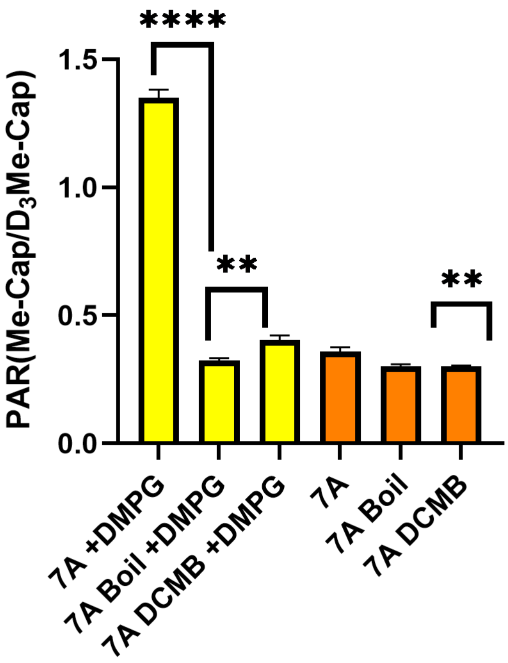


**Supplementary Figure 02.** Activity assay monitoring the formation of S-methyl captopril +/- DMPG by recombinant purified METTL7A confirms that METTL7A also requires a specific lipid environment for activity. All data are presented as the mean ±s.d.

For this captopril methylation assay, samples were incubated for 45 minutes +/- DMPG, the final DCMB concentration was 100 uM, and all other parameters and analysis techniques were performed similar to the captopril assay that is described in the in vitro 7α-thiospironolactone and captopril methylation methods section.

A

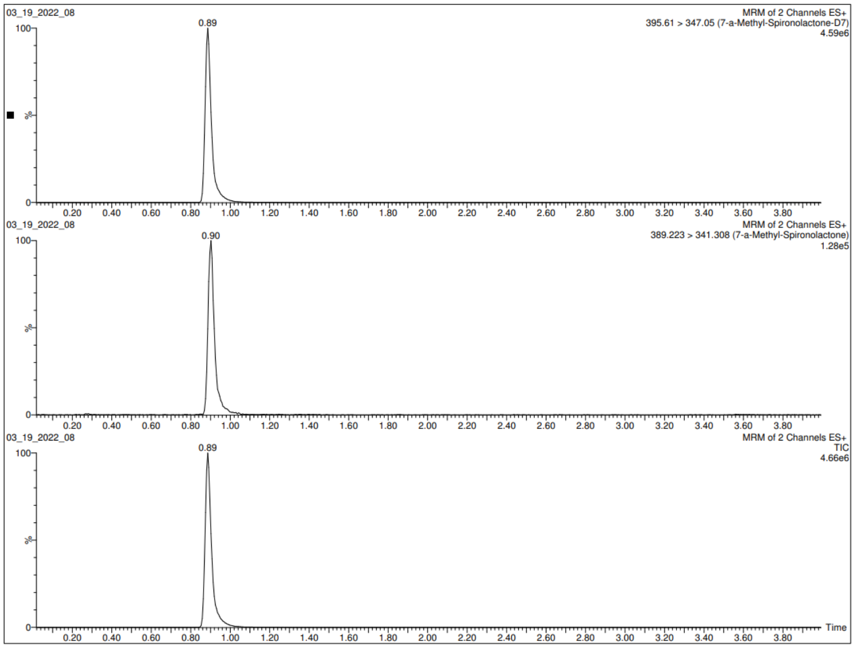


B

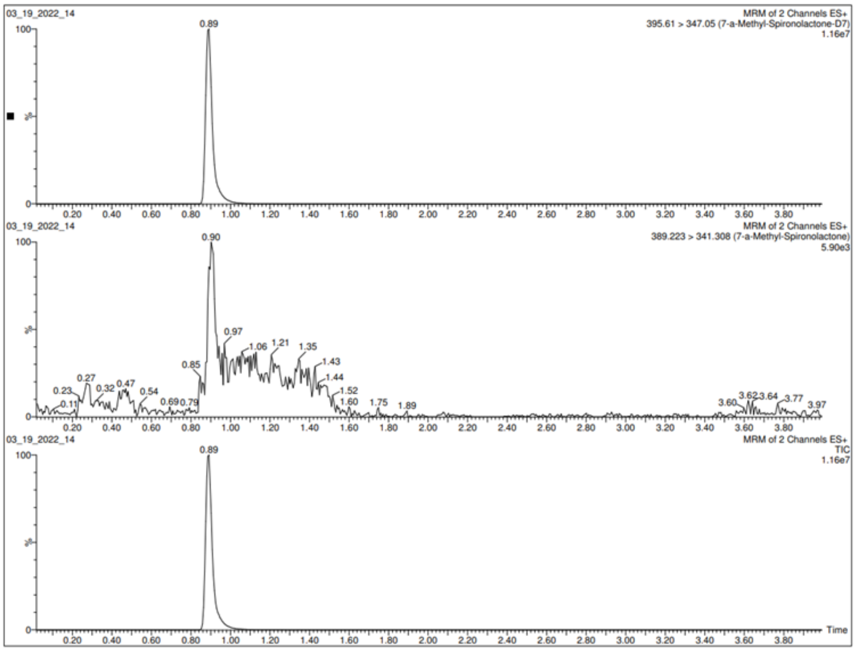


**Supplementary Figure 03.** Representative chromatogram traces from a TSL methylation activity assay with recombinant purified active (A) and boiled (B) METTL7A.

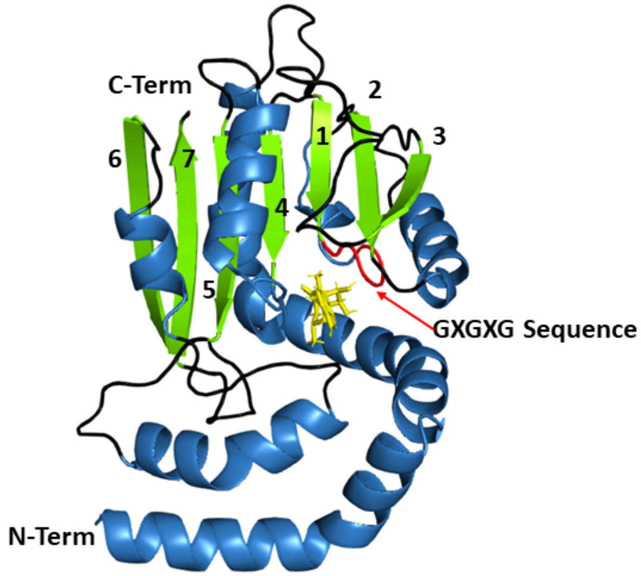


**Supplementary Figure 04.** An AlphaFold produced homology model of METTL7A with beta sheets numbered in order along protein sequence and colored green. Alpha helices are colored blue, the GXGXG sequence is colored red, and SAM is oriented in the SAM binding domain and colored yellow. The sequence of alpha helices and beta sheets along the AlphaFold-produced homology model is identical to the sequence described by Schubert et al., 2003.

The homology model for METTL7A was downloaded from The AlphaFold Protein Structure Database https://alphafold.ebi.ac.uk/files/AF-Q9H8H3-F1-model_v2.pdb. The homology model was colored using PyMOL Molecular Graphics System. Pymol renders secondary structures which allowed us to distinguish segments as alpha helices or beta sheets. SAM was docked into the METTL7A homology model with The University of Hamburg's Center for Bioinformatics protein-ligand docking software, JAMDA.

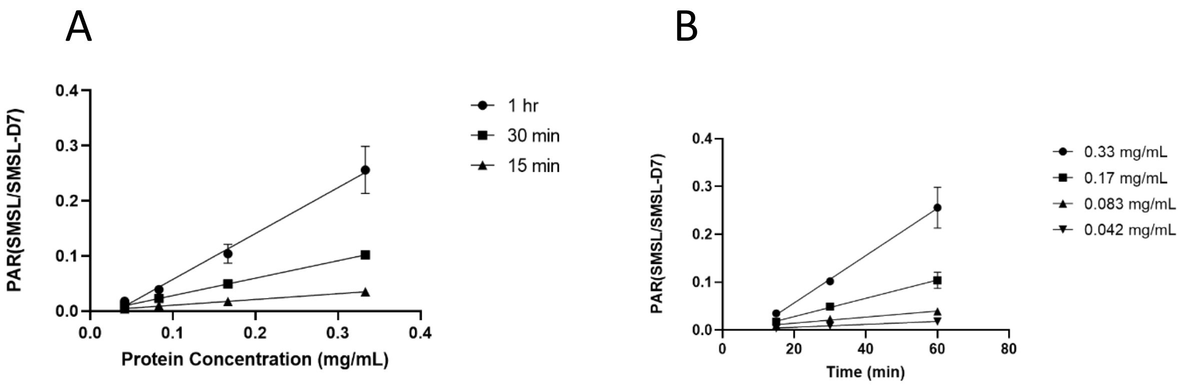


| | 1 hr | 30 min | 15 min |
|---|---|---|---|
| R squared | 0.9647 | 0.9954 | 0.9928 |

| | 0.33 mg/mL | 0.17 mg/mL | 0.083 mg/mL | 0.042 mg/mL |
|---|---|---|---|---|
| R squared | 0.9637 | 0.9613 | 0.9816 | 0.9477 |

**Supplementary Figure 05.** The methylation of TSL at various concentrations of purified recombinant METTL7A over the course of a 1 hr, 30 min, or 15 min incubation plotted with respect to protein concentration (A) or time increments (B).

Purified recombinant METTL7A was diluted to 1, 0.5, 0.25, and 0.125 mg/mL and then further diluted the purified protein into reaction buffer containing DMPG as described in the methods section; in vitro 7a-thio spironolactone and captopril methylation using recombinant METTL7A. TSL was deposited onto a 96-well plate as previously described, and the diluted protein was then added to the appropriate wells. The final protein concentrations were 0.33, 0.17, 0.083, and 0.042 mg/mL. The final TSL concentration was 100 uL. SAM was added in staggered time increments to induce the reaction. The final SAM concentration was 100 uM. The final incubation times for the various protein concentrations were 1 h, 30 min, and 15 min. The reaction was quenched, and the samples were analyzed by LC-MS/MS as was described in the method section; in vitro 7alpha-thio spironolactone and captopril methylation using recombinant METTL7A.

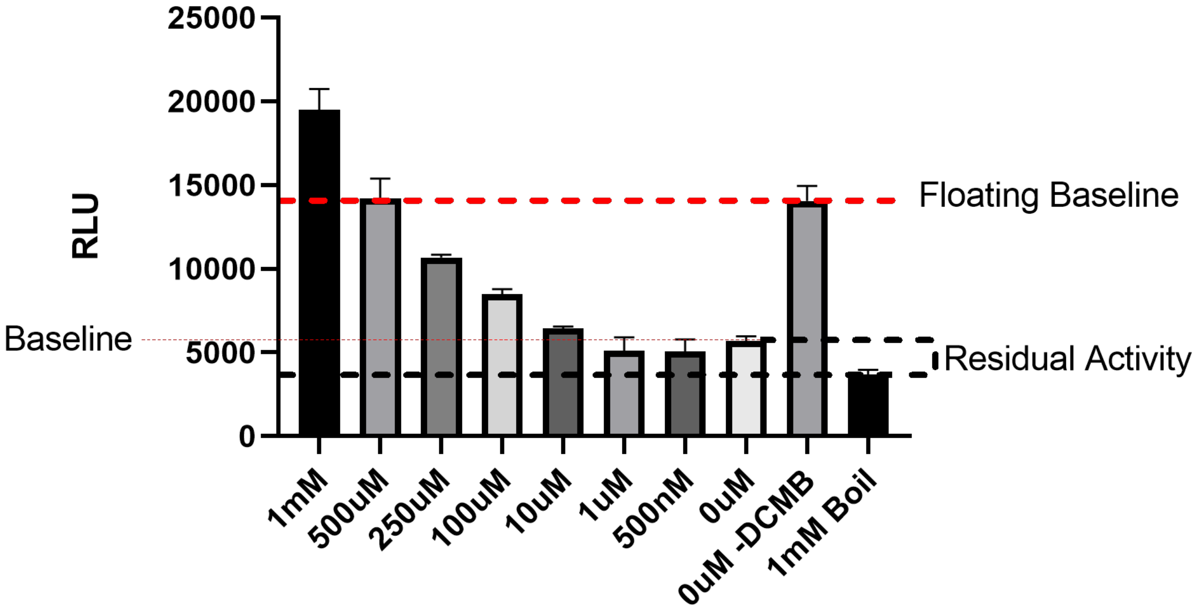


**Supplementary Figure 06.** The relative luminescence unit response from purified recombinant METTL7A incubated with various concentrations of 4-chlorothiophenol. The reaction was quenched by the addition of DCMB to a final concentration of 100 uM and all treatments were analyzed using Promega's MTaseGlo Assay. For one set of replicates, no substrate was included in the incubation and no DCMB was added to quench the reaction prior to adding the MTaseGlo development reagents. The difference between the response from this set of replicates and a set also incubated without substrate but with the addition of DCMB represents the activity that can be attributed to the methylation of a compound present in the MTaseGlo development reagents by METTL7A. There is still a response above boil in no-substrate treated replicates, but this response is low enough to identify the methylation of substrates during the activity assay.

Purified recombinant METTL7A was incubated with various concentrations of 4-chlorothiophenol for 30 minutes. The final concentration of SAM was 100 uM. All reaction conditions and analyses using the MTaseGlo Assay were identical to what was described in the substrate screening methods section. For one set of replicates, no substrate was included in the incubation and no DCMB was added to quench the reaction prior to adding the MTaseGlo development reagents.

METTL7B siRNA Sequence:

5’ CCUUCAUGUGGCAGCAAGUdTdT 3’ / 5’ ACUUGCUGCCACAUGAAGGdTdT 3’

METTL7A mRNA coding region sequence:

```
atctgtttttttcccttctgagcaatggagcttaccatctttatcctgagactggccatt
 I  C  F  F  P  F  -  A  M  E  L  T  I  F  I  L  R  L  A  I
tacatcctgacatttcccttgtacctgctgaactttctgggcttgtggagctggatatgc
 Y  I  L  T  F  P  L  Y  L  L  N  F  L  G  L  W  S  W  I  C
aaaaaatggttcccctacttcttggtgaggttcactgtgatatacaacgaacagatggca
 K  K  W  F  P  Y  F  L  V  R  F  T  V  I  Y  N  E  Q  M  A
agcaagaagcgggagctcttcagtaacctgcaggagtttgcgggcccctccgggaaactc
 S  K  K  R  E  L  F  S  N  L  Q  E  F  A  G  P  S  G  K  L
tccctgctggaagtgggctgtggcacgggggccaacttcaagttctacccacctgggtgc
 S  L  L  E  V  G  C  G  T  G  A  N  F  K  F  Y  P  P  G  C
agggtgacctgtattgaccccaaccccaactttgagaagtttttgatcaagagcattgca
 R  V  T  C  I  D  P  N  P  N  F  E  K  F  L  I  K  S  I  A
gagaaccgacacctgcagtttgagcgctttgtggtagctgccggggagaacatgcaccag
 E  N  R  H  L  Q  F  E  R  F  V  V  A  A  G  E  N  M  H  Q
gtggctgatggctctgtggatgtggtggtctgcaccctggtgctgtgctctgtgaagaac
 V  A  D  G  S  V  D  V  V  V  C  T  L  V  L  C  S  V  K  N
caggagcggattctccgcgaggtgtgcagagtgctgagaccgggagggctttctatttc
 Q  E  R  I  L  R  E  V  C  R  V  L  R  P  G  G  A  F  Y  F
atggagcatgtggcagctgagtgttcgacttggaattacttctggcaacaagtcctggat
 M  E  H  V  A  A  E  C  S  T  W  N  Y  F  W  Q  Q  V  L  D
cctgcctggcaccttctgtttgatgggtgcaacctgaccagagagagctggaaggccctg
 P  A  W  H  L  L  F  D  G  C  N  L  T  R  E  S  W  K  A  L
gagcgggccagcttctctaagctgaagctgcagcacatccaggccccactgtcctgggag
 E  R  A  S  F  S  K  L  K  L  Q  H  I  Q  A  P  L  S  W  E
ttggtgcgccctcatatctatggatatgctgtgaaatagtgtgagctggcagttaagagc
 L  V  R  P  H  I  Y  G  Y  A  V  K  -  C  E  L  A  V  K  S
```

**Supplementary Figure 07.** METTL7B siRNA sequence with region complementary to METTL7A coding region of NCBI Reference Sequence: NM_014033.4 highlighted (A). METTL7A coding region of NCBI Reference Sequence: NM_014033.4 with region complementary to METTL7B siRNA sequence highlighted B).
